# Employing Steered MD simulations for Effective Virtual Screening: Active Pharmacophore Search by Dynamic Corrections to target MKK3-MYC Interactions

**DOI:** 10.1101/2024.07.15.603633

**Authors:** Muhammet Eren Ulug, Saima Ikram, Ehsan Sayyah, Serdar Durdağı

## Abstract

The intricate relationship between mitogen-activated protein kinase 3 (MKK3) and MYC proto-oncogene protein (MYC) activation holds deep implications for the progression of cancer, particularly in the context of triple negative breast cancer (TNBC). Despite significant progress, the challenge of discovering effective MYC-targeted drugs persists, demanding innovative approaches to control MYC-dependent malignancies. A promising avenue in this pursuit involves disrupting the protein-protein interactions (PPIs) between MKK3 and MYC. The significance of this interaction is emphasized by the activation of MYC by MKK3 in diverse cell types, presenting a novel perspective for therapeutic interventions in MYC-driven pathways. In the current study, a novel *in silico* strategy to screen small molecule libraries that target the MKK3-MYC interaction was conducted. Dynamic structure-based pharmacophore models were developed and utilized for screening the small molecule libraries, enabling the identification of compounds exhibiting favorable alignment with the defined pharmacophore features. Subsequently, physics-based simulations approaches were conducted on these selected hit molecules. Steered molecular dynamics (sMD) simulations were utilized to assess the correlation between the necessary forces to dissociate candidate hit ligands from the binding pocket and their corresponding average binding free energies (MM/GBSA). Comparative analysis of the average binding free energies of the identified hits obtained from the small molecule libraries represent that the identified compounds have promising predicted binding affinities compared to the reference molecule SGI-1027. Therefore, these findings may signify a crucial advancement in our ability to control MYC activation in cancer.

## 1. Introduction

Breast cancer, a challenging global health challenge, encompasses diverse subtypes, with triple-negative breast cancer (TNBC) standing out as particularly aggressive and elusive ^1^. TNBC is characterized by the absence of estrogen receptor (ER) and progesterone receptor (PR), setting it apart from other breast cancer types and limiting the efficacy of hormone-based therapies ^2^. Unlike human epidermal growth factor receptor 2 (HER2) positive breast cancers targeted with specific anti-HER2 therapies, TNBC lacks overexpression of HER2. This unique molecular profile poses distinct challenges in treatment. The heterogeneity within TNBC, shown as a diverse array of tumors with genetic and molecular variations, further complicates treatment responses and clinical outcomes ^3,4^. The aggressive nature of TNBC is underscored by its higher likelihood of recurrence compared to other breast cancer subtypes. The absence of targeted receptors narrows treatment options, with conventional chemotherapy remaining a primary intervention ^5,6^. Despite constituting approximately 10-15% of all diagnosed breast cancers, TNBC disproportionately affects younger women with a higher risk of early recurrence and rapid metastasis ^7,8^.

The molecular landscape of TNBC is characterized by dysregulated signaling pathways, providing fertile ground for exploration. Among these pathways, the MYC oncogene emerges as a pivotal player, orchestrating crucial cellular processes and contributing to the aggressive nature of TNBC^9^. MYC activation, a hallmark of this subtype, is intricately linked to mitogen-activated protein kinase kinase 3 (MKK3), establishing a critical link in the pathogenesis of TNBC ^10^. MYC, a transcription factor, is known for its regulatory role in cell proliferation, differentiation, and apoptosis. In the context of TNBC, aberrant MYC activation contributes to uncontrolled cell growth and evasion of programmed cell death, fostering tumor progression ^11^. The dynamic interplay between MYC and MKK3 further amplifies the complexity of TNBC, as MKK3-mediated MYC activation adds layer of regulatory influence ^11^. The helix-loop-helix leucine zipper (HLH-LZ) domain of the MYC protein plays a pivotal role in its function as a transcription factor and is critical for downstream signaling in various cellular processes. The HLH-LZ domain is a structural motif found in the N-terminal region of MYC. It consists of three subdomains - helix 1 (H1), loop (L), and helix 2 (H2) - followed by a leucine zipper (LZ) motif. This domain is responsible for mediating protein-protein interactions (PPIs), including the formation of homodimers or heterodimers with other proteins.

### 1.1 Dimerization and DNA Binding

The LZ motif within the HLH-LZ domain facilitates the formation of dimers, with MYC-MAX heterodimers being a well-known example. These dimers are crucial for MYC’s ability to bind to specific DNA sequences, known as E-box elements, in the promoter regions of target genes. This binding is a key step in the transcriptional activation of downstream genes involved in cell growth, proliferation, and differentiation ^12^.

### 1.2 Transcriptional Activation

Once MYC forms a complex with its partner proteins, it acts as a transcriptional activator. The MYC-MAX heterodimer binds to E-box sequences in the promoters of target genes, recruiting transcriptional machinery and co-activators to initiate gene expression. This process leads to the upregulation of genes involved in cell cycle progression, protein synthesis, and metabolism ^13^.

### 1.3 Downstream Signaling

The activation of MYC target genes in response to its binding to the HLH-LZ domain has profound implications for downstream signaling pathways. The proteins encoded by these target genes contribute to the regulation of cell growth, proliferation, survival, and metabolism. Dysregulation of MYC activity, often observed in various cancers, can result in uncontrolled cell proliferation and contribute to tumor development. MKK3-MYC interactions and their impact on TNBC, understanding the specific role of the HLH-LZ domain is crucial. It serves as a key interface for MYC interactions with other proteins, influencing downstream signaling pathways that are implicated in the progression of TNBC ^10^.

MKK3, a key component of the MAPK signaling pathway, acts as a modulator in the MYC pathway. Its role in mediating MYC activation presents a unique opportunity for targeted interventions ^14^. By focusing on the MKK3-MYC complex, the *in silico* study holds promising potential for identifying compounds capable of modulating this interaction, offering a targeted approach for intervention in MYC-driven TNBC programs.

In the current study, we employ an innovative *in silico* method to virtually screen small molecule databases targeting the interface of the MKK3-MYC complex. This screening integrates dynamic structure-based pharmacophore modeling, similarity-based search, binary QSAR modeling, physics-based molecular simulations, steered molecular dynamics (sMD) simulations, and fragment-based design, aimed at enhancing the efficacy of the screening process.

## 2. Methods

The overall methodology of the current study is explained in Figure 1.

**Figure 1.**
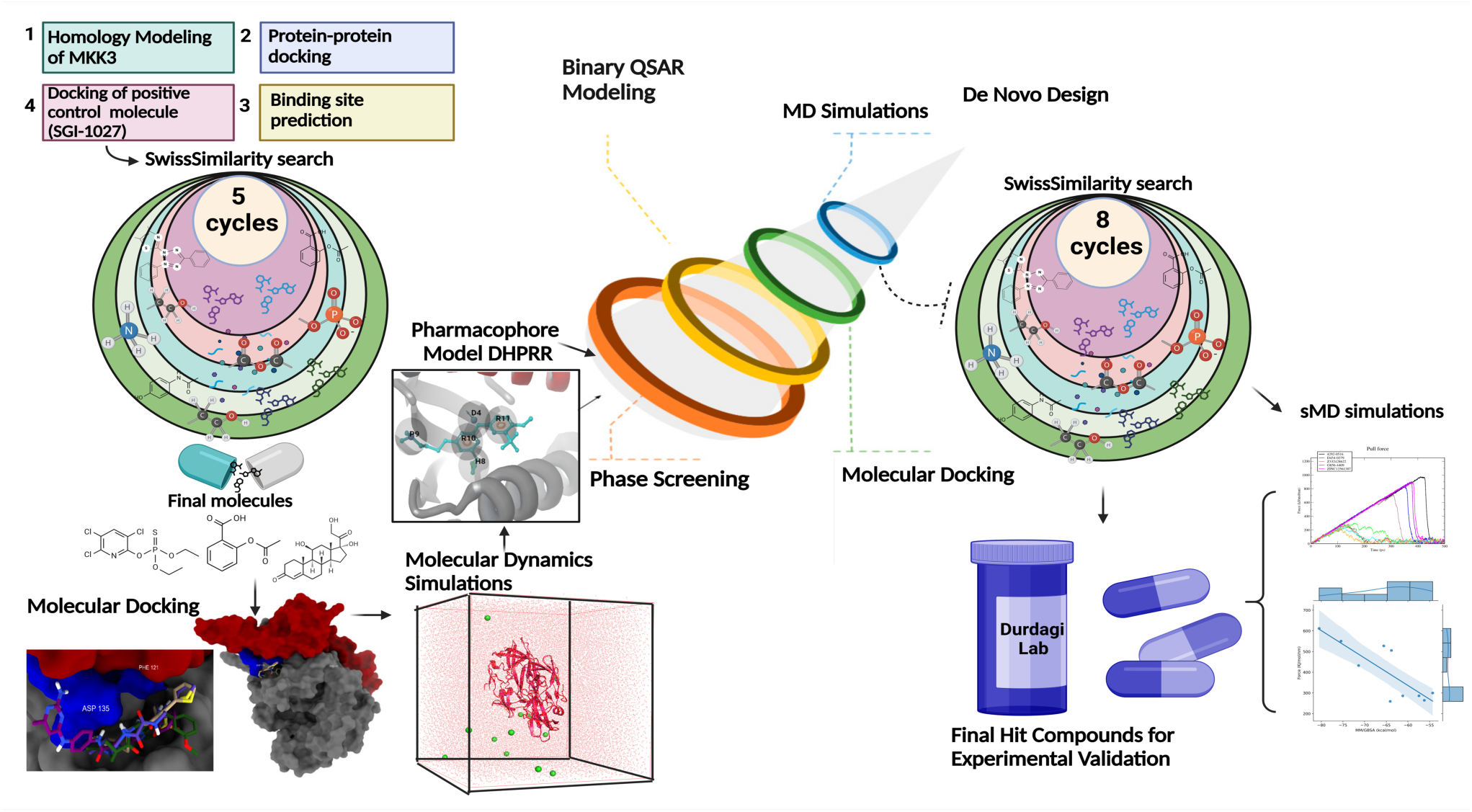
Summary of conducted methodology used in the current study.

### 2.1 Multiple Sequence Alignment

Since there is no any reported crystal structure of MKK3, we used homology modeling approach to generate the structure of MKK3. We used different programs (i.e., RosettaCM, AlphaFold2, and SwissModel) for the development of model structures for MKK3. Amino acid sequences of the mitogen-activated protein kinase (MAP2K) proteins MKK3 and MKK6, which are important in cellular signal transduction pathways, were downloaded from the UniProt server and analyzed using multiple sequence alignment (MSA) tool of Maestro ^15^. MKK3 and MKK6 show 86% sequence similarity which is significant for homology modeling. Due to their great degree of sequence similarity, the two proteins may represent homologous members of the MAP2K family and may have a conserved structure and functional role. The finding of homology between MKK3 and MKK6 has significance for comprehending the regulatory mechanisms and common functions of the MAP kinase cascade.

### 2.2 Homology Modeling: RosettaCM

The three-dimensional (3D) structural model of MKK3 was generated using the Rosetta software. (https://www.rosettacommons.org/) Rosetta, renowned for its advanced modeling capabilities, employs a high-resolution refinement protocol to predict protein structures ^16^. The primary amino acid sequence of MKK3 served as input for Rosetta, which utilized state-of-the-art algorithms to explore the conformational space and refine the model. The resulting MKK3 model was chosen for its reliability and biophysical plausibility, as assessed through various validation metrics.

### 2.3 Homology Modeling: AlphaFold2

The structural modeling of MKK3 was further complemented through the utilization of AlphaFold2, an advanced deep-learning-based method developed for predicting protein structures. AlphaFold2 employs a novel approach that combines attention-based neural networks and MSA information to generate highly accurate three-dimensional models ^17^. In our study, the primary sequence of MKK3 was used as input sequence in AlphaFold2, and the resulting model was evaluated for its structural integrity and quality. The AlphaFold-derived MKK3 structure provided an additional dimension to our computational framework, enhancing the robustness of our structural investigations.

### 2.4 Homology Modeling: SwissModel

The structural modeling of MKK3 was also accomplished using the SwissModel platform (https://swissmodel.expasy.org/), a widely recognized tool for homology modeling. SwissModel influences comparative modeling techniques to predict the three-dimensional structure of a target protein based on the experimentally determined structures of homologous proteins ^18^. The amino acid sequence of MKK3 was used in SwissModel, and the resulting model was assessed for its reliability and accuracy through various validation metrics. The SwissModel-derived MKK3 structure has been used for its robustness, served as a critical starting point for our computational investigations.

### 2.5 Protein-Protein Docking: HADDOCK2.4

The investigation of the MKK3-MYC HLH-LZ complex was further enriched through protein-protein docking simulations performed using the HADDOCK2.4 platform. HADDOCK2.4, a widely employed integrative computational docking software, is adapt at exploring PPIs by combining information from various sources, including experimental data and bioinformatics predictions ^19^. The MKK3 model obtained from SwissModel and the HLH-LZ domain of MYC were utilized as input structures for HADDOCK simulations. The HADDOCK2.4 employed a flexible and adaptive docking protocol, allowing for the exploration of conformational space and refinement of potential binding modes. The resulting docking poses were subsequently analyzed for consistency and energetically favorable conformations. The utilization of HADDOCK in current study provided valuable insights into the structural dynamics and potential interaction sites within the MKK3-MYC HLH-LZ complex.

### 2.6 Binding Pocket Search: Essential Site Scanning Analysis (ESSA)

To identify important binding sites in the target protein, a computational technique called ESSA was used ^20^. It is based on the elastic network model (ENM). These crucial locations are areas of the protein structure that are primarily responsible for defining the global mode dispersion, which is influenced by interactions between proteins and small molecules. ESSA functions by applying normal mode analysis (NMA) to the vibrational dynamics of proteins to discover locations that collectively engage large substructures of the protein at a comparatively low energetic cost. These regions are referred to as global modes or soft modes. The technique is centered on finding these critical locations that can affect the dynamics of the protein structure as a whole ^20^. ESSA incorporates data on anticipated pockets as well as their local hydrophobicity densities to identify protein binding sites. Druggable binding pocket simulations can discover potential allosteric sites, but ESSA may also effectively identify critical hot spots for high affinity binders. Additionally, the procedure is improved by including in the study particular key residues of the MKK3 and MYC proteins. Finding these crucial residues is important to accurately determine where the binding sites containing these necessary amino acid residues are located. By focusing on these residues, binding pockets relevant to these proteins can be found more quickly, which provides a tactical advantage for designing focused modulators for these interaction sites in computer-aided drug development programs.

### 2.7 Blind Docking: CB-Dock2

The blind docking program CB-Dock2 predicts binding sites for a given protein and uses a unique curvature-based cavity identification method to compute the centers and sizes (https://cadd.labshare.cn/cb-dock2/). It uses the well-known docking software AutoDock Vina to complete the docking. Blind docking is the automatic prediction of binding modes in the absence of prior knowledge about binding locations ^21^. In contrast, CB-Dock predicts the binding domains of the protein using its cavity detection technique, negating the need for users to provide predefined binding sites. This strategy uses a curvature-based method to identify holes on the protein surface, which are then ranked according to how well their size and shape complementarity with the ligand match. The centers and diameters of the top N cavities (n = 5 by default) are determined. These locations and sizes are then used in AutoDock Vina for docking along with the protein and ligand data. About the MKK3-MYC protein complex in particular, we applied this methodology to precisely locate the binding site by taking advantage of CB-Dock’s special ability to pinpoint possible interaction sites without having to be aware of the binding domains of the complex beforehand. ^21^

### 2.8. Analog Search: Swiss Similarity

SwissSimilarity, is a web tool for ligand-based virtual screening of chemical libraries to identify molecules similar to a query molecule (http://www.swisssimilarity.ch/). It provides a range of 2D and 3D molecular fingerprints that may be used to encode substances in various digital formats and measure chemical similarity. SwissSimilarity assists in identifying new compounds that may be bioactive, similar compounds with chemically distinct core structures, or readily accessible compounds for preliminary structure-activity relationship studies and is a useful tool for drug discovery and development ^22^. To initiate our study, we employed the quinoline analog SGI-1027 as a reference compound, renowned for its potency in inhibiting the MKK3-MYC interaction. This served as our starting point in identifying hit compounds with greater potency. SwissSimilarity was conducted with five cycles. In each cycle, 400 analogs of query compounds were identified and these compounds were docked to the MKK3-MYC complex. In each iteration the best-performing molecule—as indicated by the highest docking score was chosen for subsequent analysis to find more molecules that were similar. Following this procedure, a set of five potent identified compounds was assembled with the reference compound (SGI-1027).

### 2.9 Molecular Dynamics (MD) simulations

MD simulations were conducted using Desmond to unravel the dynamic behavior of the MKK3-MYC HLH-LZ domain complexes. Both short (5 ns) and long (100 ns) MD simulations were employed to capture the transient interactions and long-term stability of these complexes. The MKK3-MYC complex was placed inside a solvation box using the TIP3P water model ^23^. The complex was enclosed in an orthorhombic box with a buffer zone measuring 10 Å. The isothermal-isobaric (NPT) ensemble was used in simulations with a constant pressure of 1.01325 bar and a temperature maintained at 310 K to bring the system to thermodynamic equilibrium. First, the system was neutralized with Na^+^ ions. Next, 0.15 M of NaCl salt was added to restore the pH of the physiological condition (7.4) and the physiological ionic strength. Simulations were regulated by the Nose-Hoover thermostat and the Martyna-Tobias-Klein barostat ^24,25^. The RESPA integrator controlled the interaction forces in the simulation at different temporal granularities. The van der Waals and short-range electrostatic force calculations were performed using a 9.0 Å cutoff, and the long-range electrostatic force was calculated using the particle mesh Ewald (PME) approach ^26^ with periodic boundary constraints. The potential energy landscape of the system was determined using the OPLS3e force field ^27^. To ensure our findings, three independent replica simulations were conducted on different velocity distributions, offering insights into the reproducibility and reliability of the observed dynamics. Short MD simulations scrutinized the early stages of complex formation, while extended simulations, spanning a considerable timeframe, provided a comprehensive understanding of long-term stability and structural dynamics. The incorporation of replica simulations at different velocity distributions would further complement this analysis, offering a more thorough exploration of the energy landscape and stability profiles of the MKK3-MYC HLH-LZ domain complexes. Thus, at least three replicate simulations for each run are conducted. The analyses of trajectory data, binding free energies, and post-MD assessments contributed to unraveling the intricate molecular interplay, enhancing our understanding of the dynamic nature of the studied systems.

### 2.10 Molecular Mechanics Generalized Born Surface Area (MM/GBSA) binding free energy calculations

The MM/GBSA approach was implemented using the Prime software to calculate average binding energies ^28^. A total of 100 protein-ligand complexes were extracted at 10-frame intervals from each 1000 trajectory frames of 100 ns MD simulations. The implicit solvation model VSGB 2.0 was utilized, allowing for system-wide flexibility in the dielectric constant ^29^. The exterior of the system was modeled as a water system with a constant dielectric constant of 80, whereas the internal dielectric constant varied from 1.0 to 4.0 under the OPLS3e force field ^27^. Average values and standard deviations were found for each compound after each complex’s MM/GBSA computations.

### 2.11 Development of Dynamic Structure-Based Pharmacophore Models

Using structure-based pharmacophores in combination with all-atom MD simulations is an advanced method known as dynamic pharmacophore modeling ^30,31^. With a particular emphasis on identifying the traits that constantly predominate, our method carefully examines the pharmacophoric features that manifest throughout the simulation frames. To pick the mostly observed protein-ligand complex conformation for further research, the goal is to identify the frame with the lowest root-mean-square deviation (RMSD) based on average structure throughout the simulations ^32^. A dataset with 1000 frames was obtained from the 100 ns all-atom MD simulations of the protein-ligand interactions. Structure-based dynamic pharmacophore models were constructed using PHASE ^31^. A maximum of seven features, a minimum of 2 Å between different features, and a minimum of 4 Å between features with the same typology were used. An extra step included a receptor-specific excluded volume shell that was scaled by 0.5 to account for van der Waals radii. The protocol was improved to speed up data visualization and increase efficiency by cutting down on processing time. This optimization was made possible by an in-house bash script that used sequential commands as automation (https://github.com/DurdagiLab). The script developed by our group effectively performed a number of tasks, including identifying the ligand grid centers in each image, separating the receptor from the ligand in complexes, activating the e-pharmacophore method for in-situ docking, and producing a pharmacophore hypothesis for every frame. The outcomes were organized into categories using .csv file formats. In addition, a python script was written to sort through the hypotheses and identify the ones that were coming up most frequently during the long-span MD simulations (https://github.com/DurdagiLab). The frame with the lowest average structure RMSD was the main focus of the analysis. The ligand screening procedures were then directed by these important representative hypotheses.

### 2.12 Screening of Ligand Databases Using Dynamic Structure-Based Pharmacophore Models

Using dynamic structure-based pharmacophore model analysis as a guide, this strategy involved searching through ligand databases. Using accessible chemical conformers, the goal was to identify ligands that show a high degree of structural complementarity to the established reference pharmacophore ^33^. The fitness score of each ligand with the developed pharmacophore models at the binding pocket of the target structure is measured. The process was conducted without any restrictions and followed the uniform rejection limits that the module had established. Additionally, the screening was set up to “return at most 1 hit per molecule”, which arranged the possible hits according to their fitness scores. ∼8 million molecules from the Enamine and ChemDiv libraries were screened in this process.

### 2.13 Molecular Docking Simulations

The Glide module of Maestro molecular modeling package was used in the molecular docking procedure with extra precision (XP) ^34^ and the OPLS3e force field was configured for docking operations. The default settings of Glide/XP docking protocol were used, adding Epik ^23,33^ state penalties to the docking score. Post-docking minimization was carried out for five poses after docking, except for poses that fell inside an energy window of 0.50 kcal/mol. To increase the number of possible poses for docking, the 2.5 kcal/mol energy window was used for ring sampling. Five thousand poses for each ligand were saved during the first docking phase, and the top 800 poses were subjected to energy minimization. Expanded sample approaches were utilized to increase the sampling in the rough scoring phase, hence avoiding the exclusion of plausible positions. After finding the best poses with high docking scores the lowest-energy compounds were chosen for the next stage of the docking procedure.

### 2.14 Binary QSAR Modeling

The identification of top-docking scored molecules demonstrating chemical interactions with important residues led to their inclusion in therapeutic activity prediction protocols using the MetaCore/MetaDrug platform (https://portal.genego.com). MetaDrug employs Tanimoto Prioritization (TP) to assess the similarity between analyzed compounds and sets in binary Quantitative Structure Activity Relationships (QSAR) models, which are constructed based on diverse compounds with known activity/function on a specific protein. These models undergo rigorous testing with validation sets to determine the one with the highest specificity, sensitivity, accuracy, and Matthews Correlation Coefficient (MCC) for each therapeutic activity of interest. The prediction of therapeutic activity is derived through the ChemTree’s capacity to correlate structural descriptors to properties, utilizing the recursive partitioning algorithm. The optimal ChemTree parameters, yielding the best results, include a path length of 5, max segments of 3, a p-value threshold Bonferroni of 0.99, a p-value multiway split of 0.99, and the utilization of 50 random trees. The training set used in MetaCore/MetaDrug is composed of molecules with and without the specified property (positives and negatives, respectively) in approximately equal proportions. In instances where the number of marketed drugs exceeds 100 in the disease QSAR models, they are included; otherwise, drug candidates in clinical trials and preclinical compounds with *in vivo* activity supplement the training set. (Model Description: Potential activity against cancer. The cutoff is 0.5. Values higher than 0.5 indicate potentially active compounds. Model description: Training set N=886, Test set N=167, Sensitivity= 0.89, Specificity=0.83, Accuracy=0.86, MCC=0.72.)

### 2.15 Fragment-based Drug Design (FBDD)

To facilitate FBDD in the exploration of potential lead molecules targeting the MKK3-MYC interaction, the Auto Core Fragment in Silico Screening (ACFIS) server was employed (http://chemyang.ccnu.edu.cn/ccb/server/ACFIS2/). ACFIS, recognized as a well-established and effective online tool for FBDD, played a crucial role in generating new molecular fragments and refining lead compounds ^35^. The lead molecule, in complex with its receptor (MKK3-MYC), was used in the ACFIS, leveraging its capabilities in atomic contact fingerprints for interaction analysis. ACFIS provided a systematic exploration of chemical space, suggesting modifications and novel fragments for further consideration and optimization. The outcomes from ACFIS enriched the FBDD strategy, contributing valuable insights for the rational design of compounds with enhanced binding affinity to the MKK3-MYC interface.

### 2.16 sMD Simulations

The primary objective of the sMD simulations, conducted using the Gromacs, was to elucidate the forces necessary for ligand dissociation from the MKK3-MYC complex. CHARMM-GUI was employed for meticulous file preparation, ensuring accurate setup of the ligand-receptor system. (https://www.charmm-gui.org/). The sMD simulations were designed to apply an external force to the ligand, guiding it along predefined pathways and quantifying the pulling force required for dissociation. This analysis aimed to unravel the dynamic interactions governing ligand binding and unbinding events. To complement the sMD results, MM/GBSA calculations were performed to assess the binding free energies with required pulling forces. The integration of sMD pulling force data and MM/GBSA analysis offered a synergistic approach, providing a comprehensive perspective on the energetic and mechanical aspects of the ligand dissociation process from the MKK3-MYC complex. This dual approach facilitated a nuanced understanding of the molecular forces at play, aiding in the rational design of ligands with improved binding properties which can be adapted to use in other virtual screening efforts.

In particular, the standard Hamiltonian is extended with a harmonic time-dependent potential that acts on a particular descriptor, such as the distance between the protein and the ligand. The bound and unbound states are the two states that are transitioned between by this addition. During this transition, the system is subjected to external work, and the exerted force is calculated. Hit molecules are selected for the sMD simulations, and a constant velocity of 0.1 Å/ps is applied. In order to pull the ligand in the defined direction (i.e., negative *y*-direction) during this operation, external steering forces are applied to a reference point while maintaining the fixed center of mass of the ligand.

Using the CHARMM-GUI, the top two frames with the lowest average RMSD obtained from each replica used in standard all-atom MD simulations are recovered and prepared as starting configurations to initiate the pulling simulations ^36^. All of the components of simulated systems are subjected to parameters from the CHARMM36m force field in these simulations. The PME algorithm is used to calculate long-range electrostatics, while short-range nonbonded interactions are terminated at 1.4 nm ^37,38^. To account for the truncation of the van der Waals term, a dispersion correction is also added to the pressure and energy expressions. Periodic boundary conditions are used to simulate realistic situations, and the complicated structures are positioned inside a rectangular box with 10 Å sides on all sides. Following the minimum picture convention, this arrangement leaves plenty of room for pulling simulations along the *y*-axis. The GROMACS package, v.2022, is used for the actual pulling simulations, making use of its newly added capabilities and improved pull code. First, the steepest descent algorithm is used for a minimization step. Then, NPT ensemble with weak coupling is carried out for 500 ps. Following equilibration, the constraints on ligands are released, but they remain on the protein to provide an immobile reference for the pulling simulations. An ultimate center-of-mass distance of roughly 5.5 nm is reached between the ligand and protein by applying a spring constant of 100 kJ/(mol·nm^2^) and maintaining a constant rate of 0.1 Å/ps during the ligand pulling phase.

## 3. Results and Discussion

Exploring the dynamic interactions between MKK3 and MYC is crucial in decoding the molecular intricacies underlying TNBC. MKK3, a kinase known for its involvement in stress-activated protein kinase pathways, has emerged as a key player in the intricate network of signaling cascades within cancer cells. Concurrently, MYC, a transcription factor notorious for its potent oncogenic capabilities, is implicated in the pathogenesis of various cancers, including TNBC. The convergence of these two entities in a molecular alliance underscores the significance of the MKK3-MYC complex in TNBC, where aberrant activation of MYC is a hallmark feature.

### 3.1 Homology Modeling

Understanding the regulatory mechanisms and shared roles of the MAP kinase cascade depends on the finding of homology between MKK3 and MKK6 (Figure S1). In the absence of a crystal structure for MKK3, we employed computational tools such as SwissModel, Rosetta, and AlphaFold to construct a reliable structural model of MKK3 (Figure S2). The choice of the SwissModel structure for MKK3 was driven by several critical factors that ensure the model’s reliability and suitability for further analysis. Firstly, SwissModel has established itself as a widely recognized and reliable platform for homology modeling, renowned for its accuracy and efficiency in generating high-quality structural models based on known template structures. Additionally, SwissModel employs advanced algorithms and computational methodologies that incorporate various parameters such as sequence alignment, structural similarity, and template quality assessment to produce refined and biophysically plausible models. Furthermore, our decision to prioritize the SwissModel-generated MKK3 structure was influenced by the comprehensive evaluation of multiple computational tools, including Rosetta and AlphaFold. While these tools are valuable resources in protein structure prediction, SwissModel consistently outperformed others in terms of providing a structurally sound and validated model for MKK3 based on quality metrics and validation parameters, ensuring the fidelity and reliability of the resulting model. Moreover, utilizing the SwissModel-derived MKK3 structure aligns with the MKK6 (sequence similarity 87%) rigorous standards of structural biology research, where the selection of the most accurate and biophysically relevant model is dominant. By analyzing all these parameters, we could confidently proceed with our investigation into the regulatory mechanisms and shared roles of the MAP kinase cascade, facilitating a deeper understanding of MKK3 function and its implications in cellular signaling pathways. The crystal structure of MYC was downloaded from the protein data bank (PDB), ID: (1NKP). Only the HLH-LZ domain of the MYC was considered for further study.

### 3.2 Protein-protein Docking

After obtaining the MKK3 model through SwissModel and selecting the HLH-LZ domain of the MYC protein, protein-protein docking simulations using the HADDOCK2.4 platform were conducted. The rationale behind this approach was to explore the potential structural interactions and binding modes between MKK3 and the essential functional domain of MYC implicated in downstream signaling. Residues Phe121-Asp135 of MKK3 are involved in PPI. The docking results revealed a series of putative binding poses, each characterized by a distinct orientation of the MKK3-MYC HLH-LZ complex. Clustering analysis was performed to identify the most prevalent and energetically favorable conformations (Figure 2, left).

**Figure 2.**
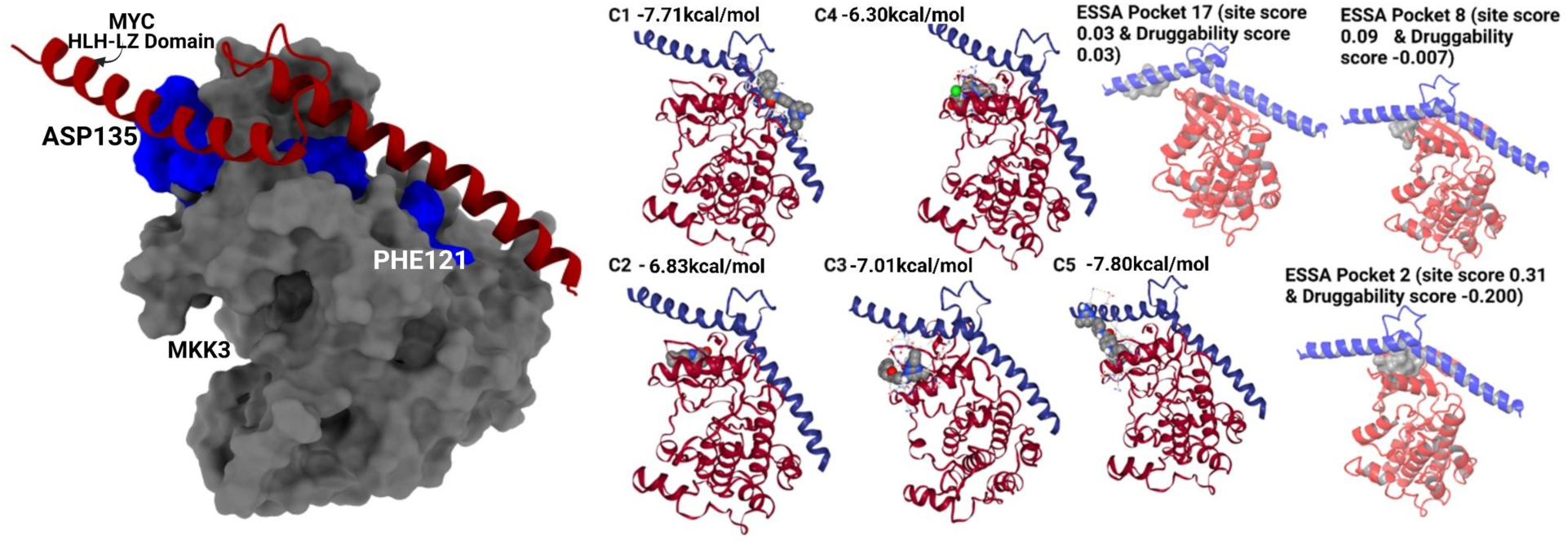
(left) Protein-protein docking complex of MKK3 and HLH-LZ domain of MYC. The red color indicates the HLH-LZ domain, the grey color shows MKK3 and the blue color shows interacting residues Phe121-Asp135 of MKK3 with MYC. (right) Comparison of binding site poses: ESSA software and CB-Dock2-generated binding site poses (C1-C5) depict the predicted binding poses within the target binding site.

The interface analysis indicated that residues involved in the interaction were predominantly located in the helical regions of MKK3 and the HLH-LZ domain of MYC. This observation aligns with the known structural motifs responsible for mediating PPIs, reinforcing the biological relevance of our docking results ^10^. Additionally, the binding poses suggested potential key residues contributing to the stability of the MKK3-MYC complex, laying the foundation for further mutagenesis studies to validate the predicted interactions.

### 3.3 Validation of Binding Site

After identifying the putative binding site residues from the literature ^10^, we aimed to validate our docking approach through blind docking simulations using CB-Dock2 and ESSA to assess the ensemble-based similarity scores of the generated docking poses. SGI-1027 is the well-known inhibitor of the MKK3-MYC PPI ^10^. We used SGI-1027 as a reference molecule for our study and for the validation of the binding pocket. CB-Dock2 was employed for blind docking, where the binding site information is not explicitly provided to the algorithm, allowing us to assess the robustness and reliability of our docking predictions ^39,40^. The blind docking simulations with CB-Dock2 yielded a range of binding poses, enabling us to evaluate the accuracy of our methodology in predicting the binding site and orientation of the MKK3-MYC HLH-LZ complex. Analysis of the blind docking results indicated a convergence of predicted binding poses within the known binding site region, providing validation for our docking protocol and supporting the credibility of our predicted binding interactions (Figure 2, right).

To complement the blind docking validation, we employed the ESSA software to assess the ensemble-based similarity scores of the generated docking poses. ESSA offered a valuable tool for evaluating the consistency and reliability of docking results by comparing the ensemble of predicted structures ^20^. The analysis revealed a high degree of convergence among the top-scoring docking poses, further validating the accuracy and precision of our approach in capturing the energetically favorable conformations of the MKK3-MYC HLH-LZ complex. Additionally, we compared the binding modes and interacting residues identified through blind docking with CB-Dock2 and ESSA analysis with the literature-derived binding site information. The congruence observed between our predictions and the known binding site residues provides additional confidence in the reliability of our computational modeling strategy.

In summary, the blind docking simulations with CB-Dock2 and subsequent ESSA served as robust validation steps, confirming the accuracy of our approach in predicting the binding modes and key interacting residues within the MKK3-MYC HLH-LZ complex. These results reinforce the credibility of our computational model and lay the groundwork for the subsequent exploration of small-molecule modulators, with SGI-1027 as the reference, in targeting this critical interaction.

### 3.4 SwissSimilarity Screening and Iterative Refinement

SwissSimilarity, a powerful cheminformatics tool, was employed to perform an iterative screening process aimed at identifying potential lead compounds against the target of interest. In the initial cycle, we used a known inhibitor SGI-1027 in SwissSimilarity, which generated a diverse set of 400 molecules sharing structural similarities. These molecules were then used in the molecular docking tool Glide XP, and the top-ranked compound was selected for the subsequent cycle. This process was iteratively repeated for a total of 5 cycles. Figure 3 represents the binding pose of SGI-1027 analog that was used for pharmacophore screening.

**Figure 3.**
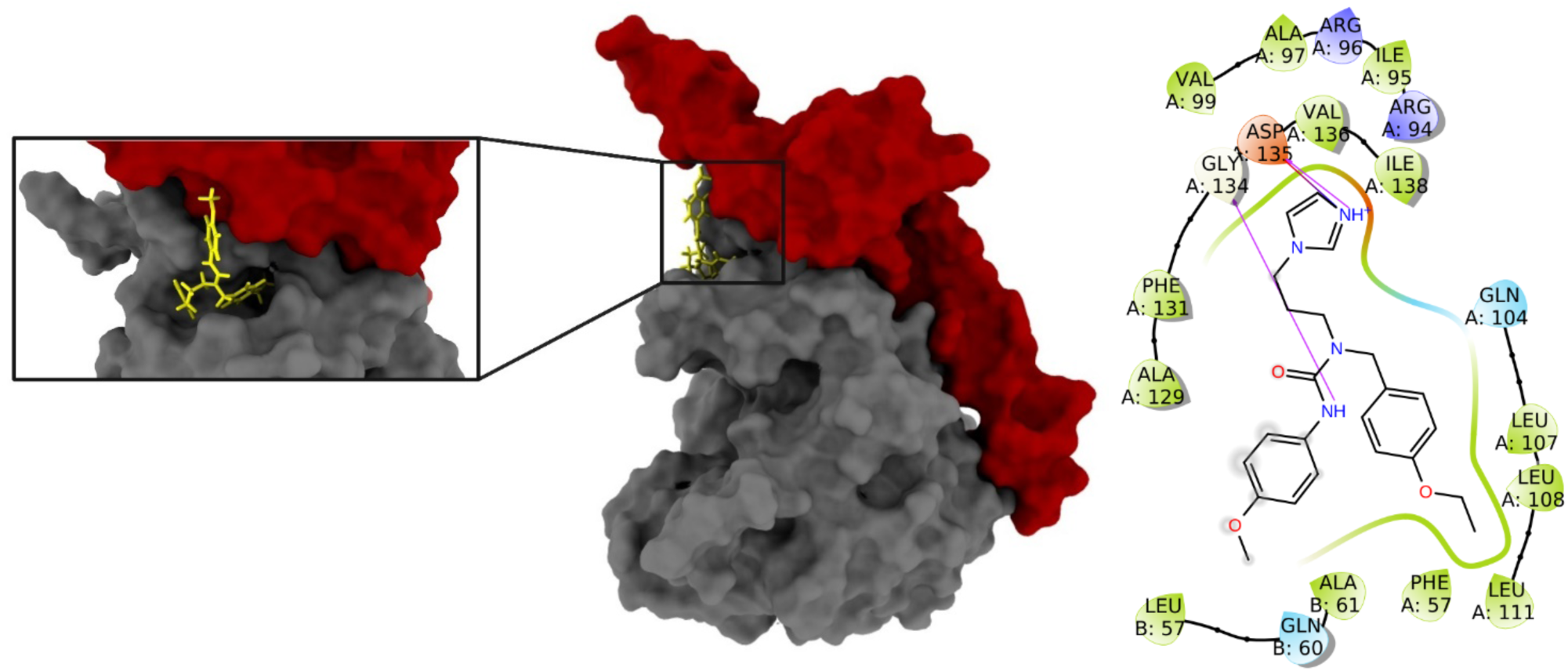
Surface representation of MKK3-MYC protein-protein interaction complex. The red and gray colors represent MYC and MKK3 structures, respectively. The figure shows the binding site and top-docking pose of the hit compound that was used for grid generation and pharmacophore screening. Additionally, a 2D interaction map is included, illustrating the detailed interactions between the MKK3-MYC complex and the hit compound (Z332428622).

SGI-1027 has docking score of -5.33 kcal/mol and its ligand efficiency score (i.e., docking score per nonhydrogen atoms) was measured as -0.15 kcal/mol. The first two cycles of similarity search yielded suboptimal results, with the top-ranked molecules exhibiting similar docking scores with SGI-1027. (Tables S1 and S2) However, in the third cycle, a notable improvement was observed, as the selected molecules demonstrated enhanced predicted binding affinity (Table 1). This observed enhancement in docking scores suggests a refinement in the screening process, where the iterative nature of SwissSimilarity allowed for the exploration of chemical space and the identification of more promising lead candidates. The subsequent cycles further capitalized on this improved trajectory, ultimately leading to the identification of a top-ranking compound with superior binding characteristics.

**Table 1.**
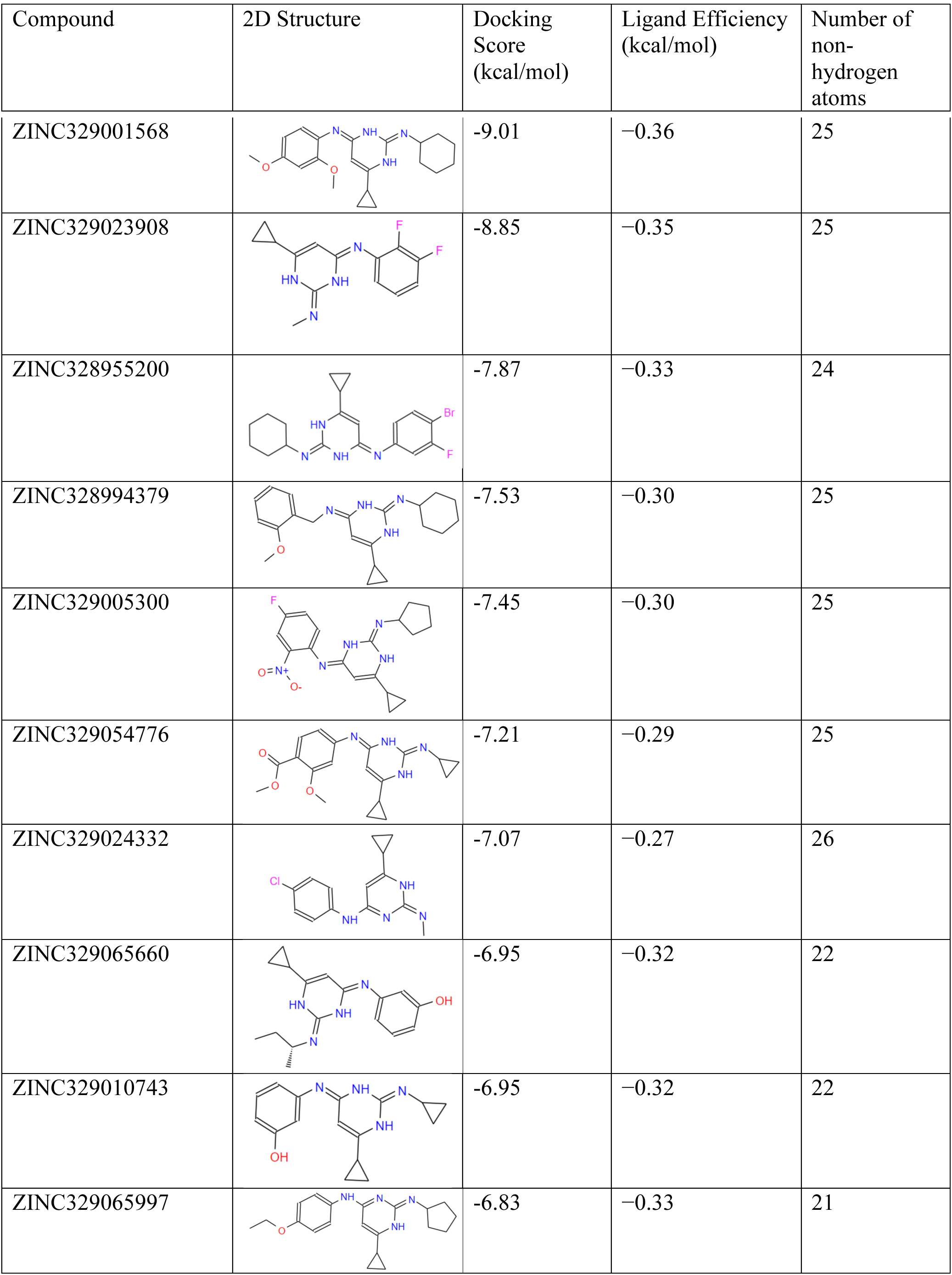
ZINC329065661 (-6.95 kcal/mol) was used for Swiss similarity 3^rd^ round. Table shows the analogs of ZINC329065661 along with docking and ligand efficiency scores (i.e., docking score per number of non-hydrogen atoms).

| Compound | 2D Structure | Docking Score (kcal/mol) | Ligand Efficiency (kcal/mol) | Number of non-hydrogen atoms |
| --- | --- | --- | --- | --- |
| ZINC329001568 |  | -9.01 | -0.36 | 25 |
| ZINC329023908 |  | -8.85 | -0.35 | 25 |
| ZINC328955200 |  | -7.87 | -0.33 | 24 |
| ZINC328994379 |  | -7.53 | -0.30 | 25 |
| ZINC329005300 |  | -7.45 | -0.30 | 25 |
| ZINC329054776 |  | -7.21 | -0.29 | 25 |
| ZINC329024332 |  | -7.07 | -0.27 | 26 |
| ZINC329065660 |  | -6.95 | -0.32 | 22 |
| ZINC329010743 |  | -6.95 | -0.32 | 22 |
| ZINC329065997 |  | -6.83 | -0.33 | 21 |

The success of the iterative SwissSimilarity approach underscores its utility in navigating chemical diversity and refining the selection process over multiple cycles. This methodology not only highlights the importance of persistence in virtual screening but also emphasizes the dynamic nature of structure-based drug design, where iterative refinement plays a pivotal role in uncovering high-affinity ligands against the target of interest (Supplementary Tables S3 and S4).

### 3.5 MD Simulations of Top-Ranked Molecules

The top-5 compounds based on docking scores obtained from each cycle of the SwissSimilarity search were subjected to MD simulations using the Desmond for both short (5 ns) MD and long (100 ns) MD simulations with three replicates. The MD parameters, including docking scores, ligand efficiency, and MM/GBSA scores, were precisely assessed to evaluate the predicted binding affinity and efficiency of each compound (Table 2).

**Table 2.**
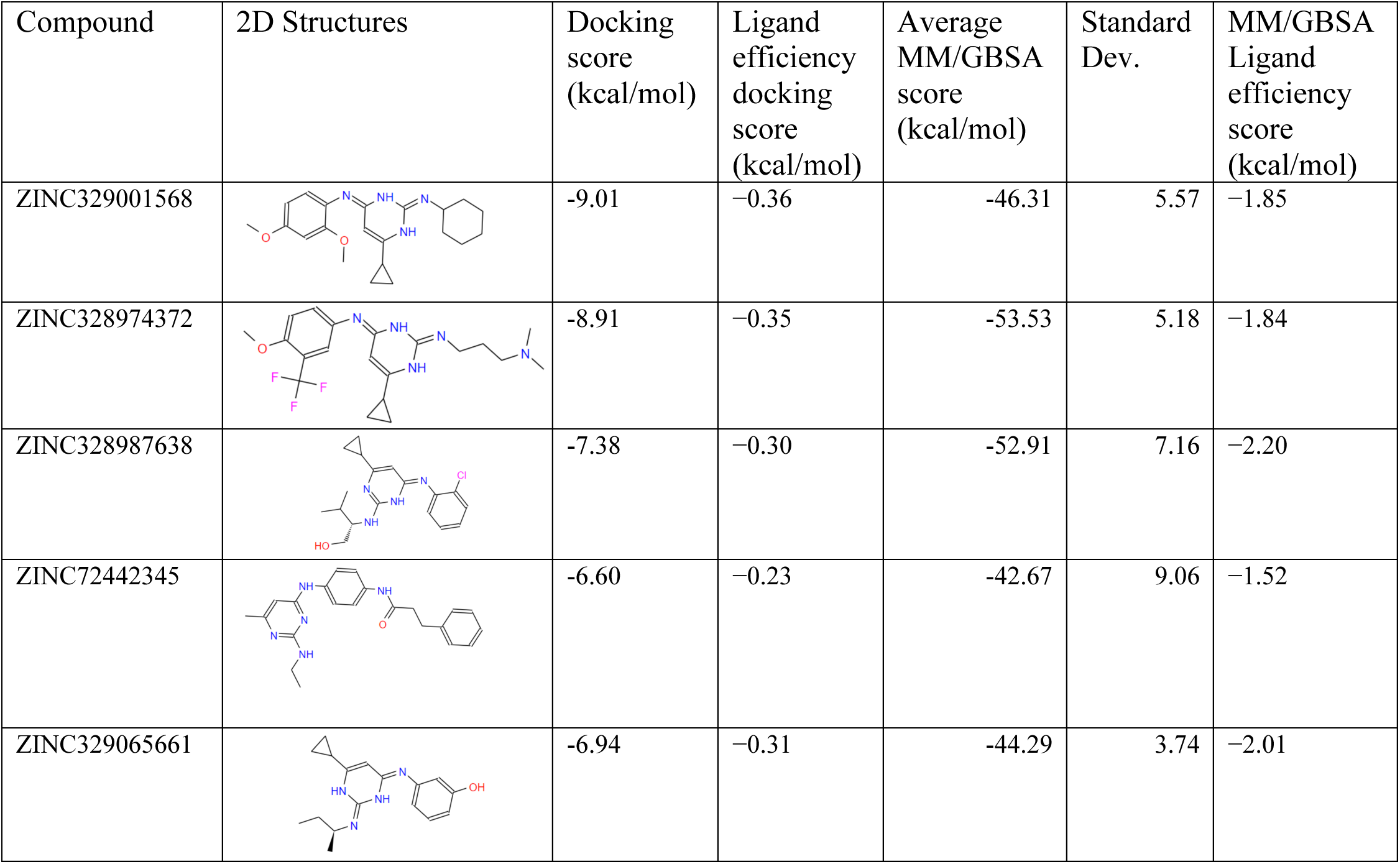
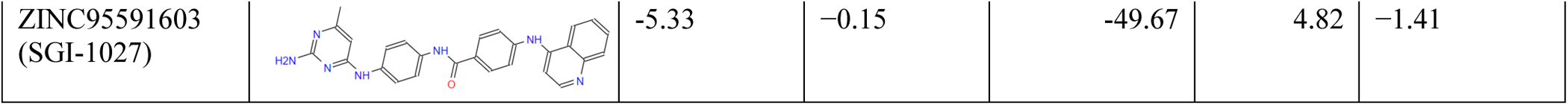
2D structure, docking score, ligand efficiency and average MM/GBSA scores of top-5 molecules obtained from each round of SwissSimilarity.

| Compound | 2D Structures | Docking score (kcal/mol) | Ligand efficiency docking score (kcal/mol) | Average MM/GBSA score (kcal/mol) | Standard Dev. | MM/GBSA Ligand efficiency score (kcal/mol) |
| --- | --- | --- | --- | --- | --- | --- |
| ZINC329001568 |  | -9.01 | -0.36 | -46.31 | 5.57 | -1.85 |
| ZINC328974372 |  | -8.91 | -0.35 | -53.53 | 5.18 | -1.84 |
| ZINC328987638 |  | -7.38 | -0.30 | -52.91 | 7.16 | -2.20 |
| ZINC72442345 |  | -6.60 | -0.23 | -42.67 | 9.06 | -1.52 |
| ZINC329065661 |  | -6.94 | -0.31 | -44.29 | 3.74 | -2.01 |
| ZINC95591603<br>(SGI-1027) |  | -5.33 | -0.15 | -49.67 | 4.82 | -1.41 |

The results reveal significant trends across the top-ranked molecules from different SwissSimilarity cycles. Interestingly, the dynamics observed during the MD simulations provided additional context to the binding characteristics, shedding light on the stability and adaptability of these compounds within the binding pocket. Reference molecule SGI-1027 was included in the analysis to benchmark and contextualize the performance of the selected candidates. The MD simulation results underscore the dynamic nature of ligand-protein interactions, revealing distinct behaviors among the top-ranked molecules. The compounds exhibiting favorable MD profiles, as indicated by improved ligand efficiency and average MM/GBSA scores, merit further consideration for their potential as lead candidates.

### 3.6 Dynamic Structure-Based Pharmacophore Modeling

Pharmacophore hypotheses were generated from the MD trajectories of the top-5 molecules obtained from each SwissSimilarity cycle together with SGI-1027, capturing the essential features critical for ligand binding. During the 100 ns MD simulations, a comprehensive analysis was conducted to identify crucial features within the trajectories associated with the ligand-protein complex. The focus was on explaining features that contribute significantly to the stability and interactions within the binding site. Table 3 represents the most frequently viewed pharmacophore features in the trajectories of selected molecules.

**Table 3.** Common features in dynamic structure-based pharmacophore modeling of SGI-1027 and top 5 molecules obtained from each round of SwissSimilarity. Pharmacophore features: A, hydrogen-bonding acceptor; D, hydrogen-bond donor; R, aromatic ring; H, hydrophobic.

| Compound | Common Pharmacophore Features | Observed pharmacophore features (%) | Representative frame # | RMSD of Representative Frame (Å) |
| --- | --- | --- | --- | --- |
| SGI-1027 | ARRRR | 20.2 | 589 | 1.25 |
| ZINC72442345 | DRRR | 43.1 | 482 | 0.85 |
| ZINC328974372 | DHPRR | 20.9 | 656 | 0.84 |
| ZINC328991568 | HPRR | 54.9 | 405 | 0.86 |
| ZINC329065661 | AHRR | 64.7 | 605 | 0.90 |
| ZINC328987638 | DHRR | 50.2 | 745 | 1.03 |

Within the representative frames, frame number #656 of ZINC328974372 shows the lowest RMSD (0.84 Å) to the average protein structure throughout the simulations as compared to the representative frame of SGI-1027 (Table 3), and average MM/GBSA score of this lead molecule has a better score than SGI-1027 (Table 2). Based on this analysis, a subset of features, referred to as DHPRR features, emerged as particularly prominent and recurrent and decided for further screening (Figure 4).

**Figure 4.**
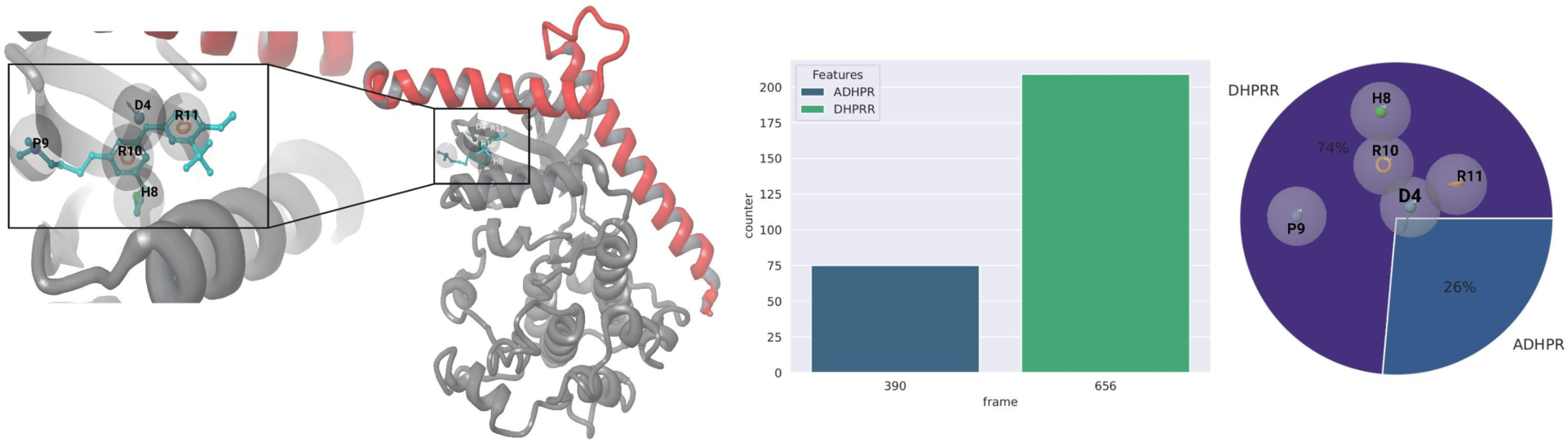
(left) Binding pose of reference ligand in frame #656 that best aligned with the hypothesis. (rigth) Representation of the MD trajectories highlighting the hypothesis and its associated DHPRR features. The DHPRR features, depicted with distinct markers, are consistently present throughout the trajectories, as illustrated in the plot. This figure provides a dynamic snapshot, linking the feature hypothesis to specific frames and emphasizing its recurrent nature across the MD simulations.

These features demonstrated a consistent presence throughout the trajectory, indicating their persistent and potentially crucial role in the ligand-protein interactions with the RMSD of 0.84 Å. Throughout the simulations, a total of 209 trajectories, out of the 1000 collected frames, exhibited DHPRR features. This high frequency across the trajectories underscores the reproducibility and prevalence of these specific features, emphasizing their importance in the context of the ligand-protein binding dynamics (Figure 4). The presence of DHPRR features in a substantial portion of the MD trajectories suggests their involvement in maintaining the stability of the ligand-protein complex. These features may contribute to critical interactions, structural conformations, or other factors that influence the overall binding affinity. This method enhances the understanding of molecular recognition, aiding in the design of novel and effective therapeutic agents. The trajectories of six ligands were used to generate a pharmacophore model (Figure S3).

Pharmacophore hypotheses were generated between 3 to 7 pharmacophore features, but, the 5-sited hypothesis (DHPRR, D: hydrogen-bond donor, P: positively charged, H: hydrophobic, R: aromatic) is represented to occur more frequently compared to other features during the MD simulations. To determine the most frequently observed receptor-ligand-derived pharmacophore hypothesis in each trajectory, frames displaying these features were ranked according to their RMSD values in comparison to the average protein structure of the respective MD simulations. The selected optimized representative structures were then employed for grid generation and virtual screening. The pharmacophore model was then employed to conduct virtual screening against the ChemDiv 1.6 million (https://www.chemdiv.com/) and Enamine (https://enamine.net) (around 400.000) small molecule libraries. Among more than 2 million compounds in total, 16766 compounds showed 5 out of 5 features that fit with the model. 16766 compounds were then used in molecular docking simulations and top-docking scored 100 compounds are identified. These top-100 hits are used for further evaluation (Figure 5, left).

**Figure 5.**
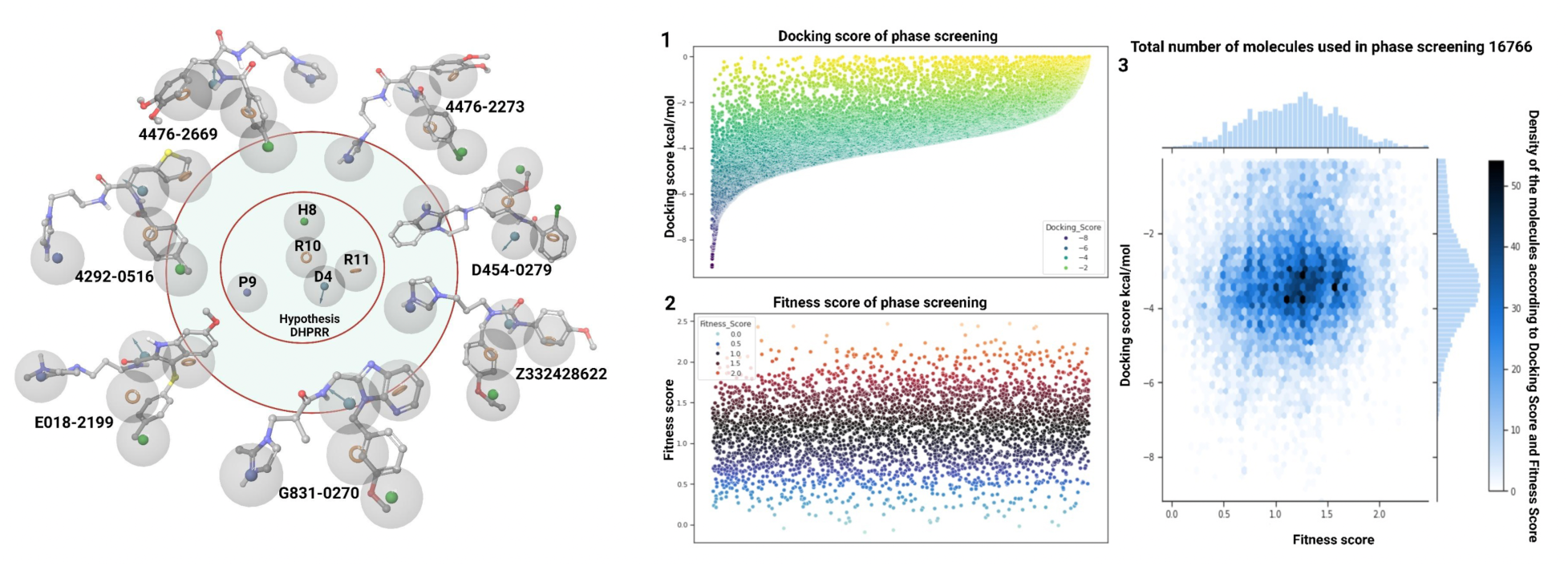
**(left)** Top-screened molecules that fit all 5 features of the DHPRR hypothesis during phase screening among 16766. **(right)** Comparative analysis of Glide/XP docking and fitness scores for 16766 compounds in DHPRR hypothesis. These 3 graphs illustrate the docking scores, fitness scores, and their correlation. Graph 1 depicts individual XP docking scores, Graph 2 shows fitness scores and Graph 3 provides a visual representation of the relationship between docking and fitness scores.

The utilization of MD trajectories for pharmacophore generation offered a dynamic perspective on ligand binding, capturing the inherent flexibility and adaptability of the ligands within the binding pocket. The subsequent virtual screening against these pharmacophores from diverse chemical databases provided a targeted approach to identify compounds with a high likelihood of mimicking the essential interactions observed in the MD simulations. This integrative strategy, combining structural insights from MD with virtual screening, enhances the probability of selecting lead candidates with a robust binding profile for further experimental validation. The generated pharmacophore hypotheses, along with the selected compounds, represent a strategic advancement in the rational design of potential therapeutics targeting the MKK3-MYC.

### 3.7 Docking and MD simulations of identified compounds from pharmacophore screening

A comprehensive virtual screening approach was employed to evaluate the predicted binding affinities of 16766 compounds obtained from ChemDiv and Enamine libraries. The Glide/XP docking algorithm, was employed to meticulously rank these compounds based on their potential to form stable complexes with the target protein. Molecular docking analysis of lead compounds shows notable predicted binding affinities and interaction profiles against the MKK3-MYC complex (Figure 5, right).

Among them, 4292-0516 exhibited substantial interactions with crucial residues at the binding pocket with a docking score of -8.48 kcal/mol. This compound demonstrated a hydrogen bond and salt bridge interaction with the crucial residue Asp135, while engaging in hydrophobic interactions with Phe131, Leu107, Leu108, Ala129, Leu111, and Phe57 (Figure 6).

**Figure 6.**
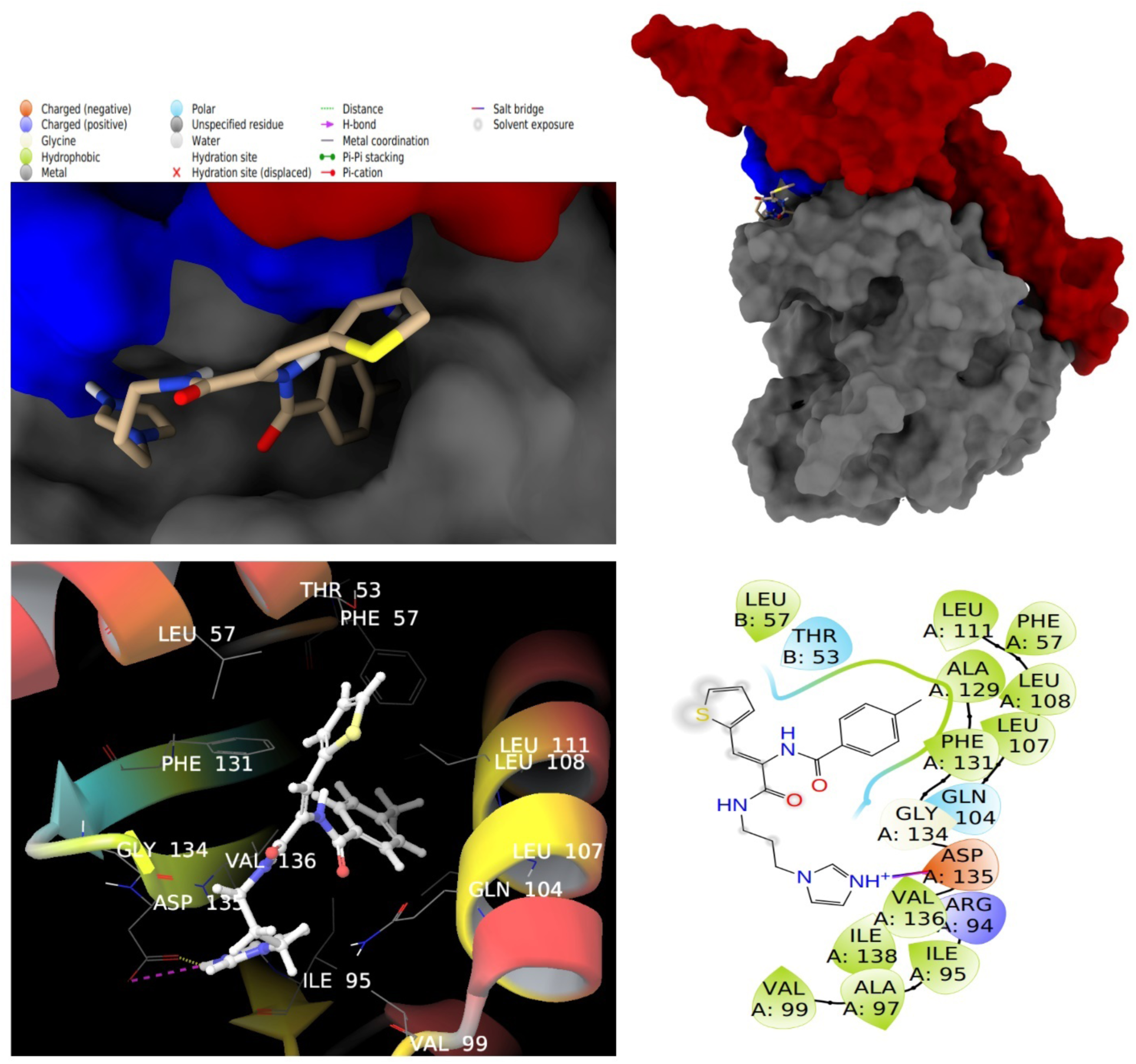
3D and 2D interaction map of compound 4292-0516. The blue color shows the interface region of MKK3 and MYC HLH-LZ.

Similarly, compounds 4476-2273 and 4476-2669 exhibited analogous interactions with Asp135, securing docking scores of -8.18 kcal/mol and -8.03 kcal/mol, respectively. (Table 4) The former achieved a fitness score of 1.1, while the latter obtained a fitness score of 0.72. These findings highlight the consistent involvement of Asp135 in ligand binding, emphasizing its significance in the MKK3-MYC interaction network. Furthermore, C090-0364 shows a docking score of -7.69 kcal/mol and a fitness score of 1.24, primarily through hydrophobic interactions. (Table 4) Compound D454-0279, with a docking score of -6.76 kcal/mol and a fitness score of 1.26,

**Table 4.**
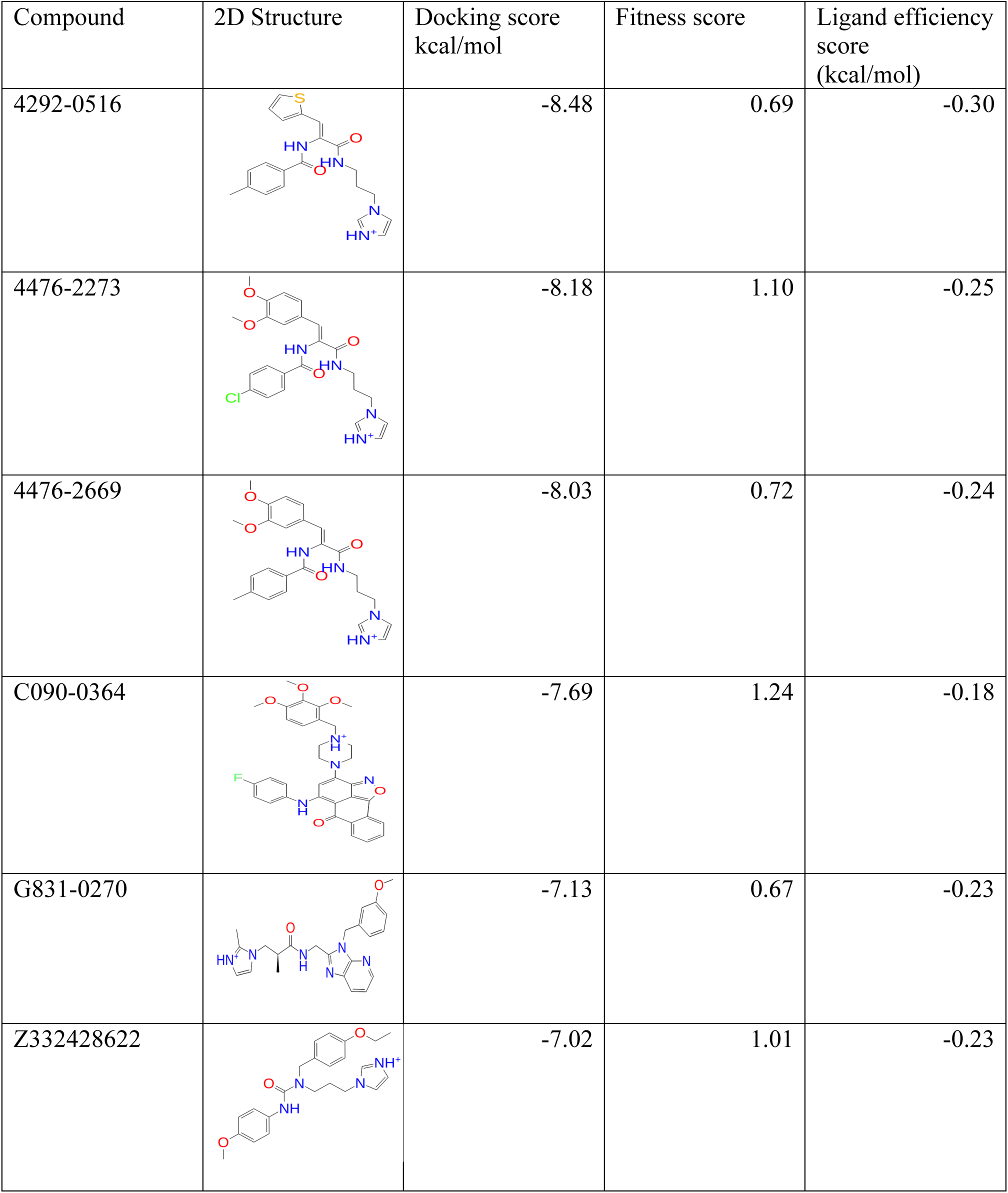
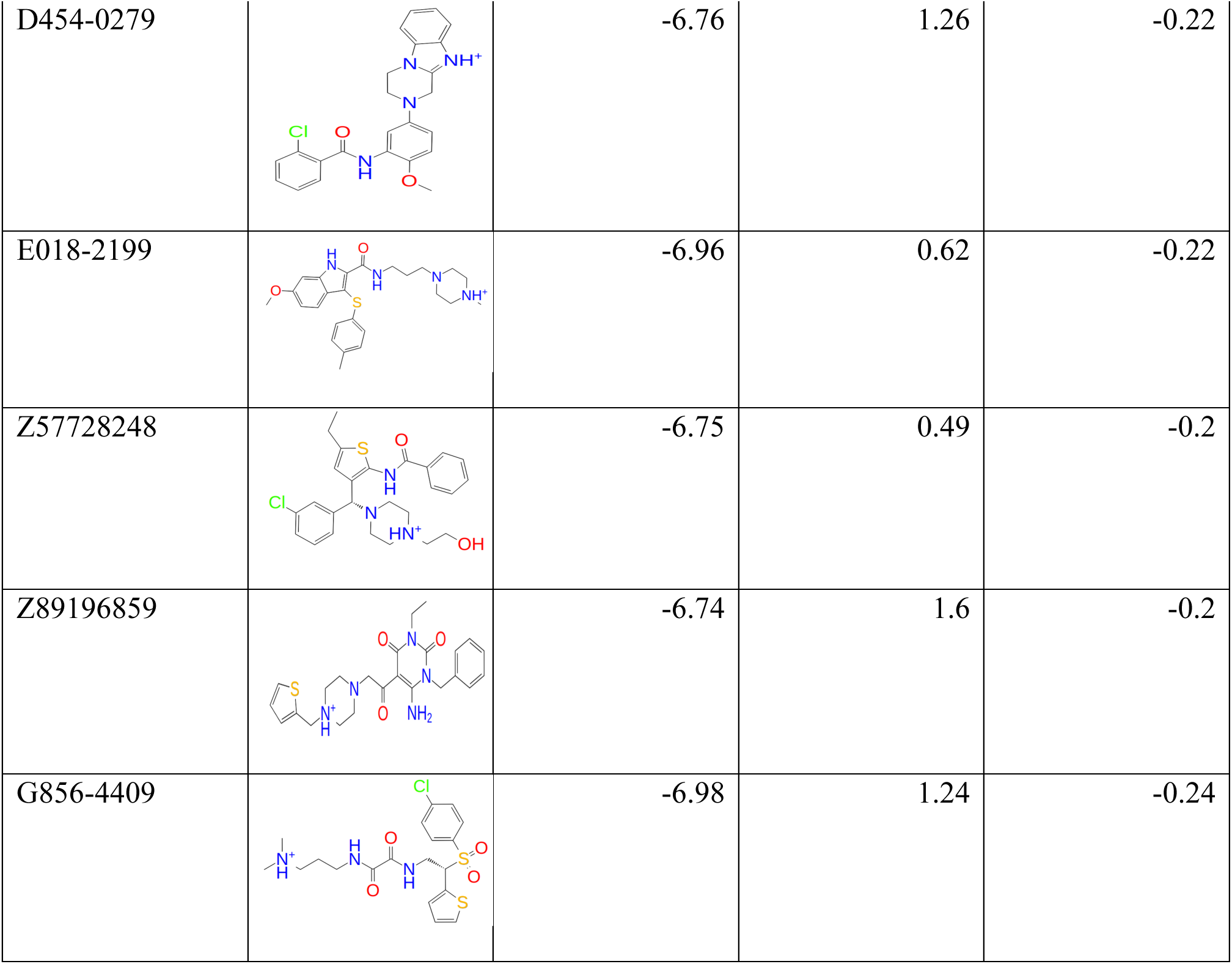
2D structure, docking score, fitness score, and ligand efficiency of top-hit candidate molecules obtained from pharmacophore feature screening.

| Compound | 2D Structure | Docking score<br>kcal/mol | Fitness score | Ligand efficiency<br>score<br>(kcal/mol) |
| --- | --- | --- | --- | --- |
| 4292-0516 |  | -8.48 | 0.69 | -0.30 |
| 4476-2273 |  | -8.18 | 1.10 | -0.25 |
| 4476-2669 |  | -8.03 | 0.72 | -0.24 |
| C090-0364 |  | -7.69 | 1.24 | -0.18 |
| G831-0270 |  | -7.13 | 0.67 | -0.23 |
| Z332428622 |  | -7.02 | 1.01 | -0.23 |
| D454-0279 |  | -6.76 | 1.26 | -0.22 |
| E018-2199 |  | -6.96 | 0.62 | -0.22 |
| Z57728248 |  | -6.75 | 0.49 | -0.2 |
| Z89196859 |  | -6.74 | 1.6 | -0.2 |
| G856-4409 |  | -6.98 | 1.24 | -0.24 |

maintained crucial salt bridge and hydrogen bonding interactions with Asp135, representing its potential as a binding modulator. (Table 4) In comparison, the reference molecule SGI-1027, while exhibiting a docking score of -5.35 kcal/mol, formed three hydrogen bonds with Gln64, Glu133, and Gly134. While these interactions are significant, the docking scores of the newly identified hit candidate compounds surpassed that of the reference, signifying their potential as stronger binders to the MKK3-MYC complex (Figure 7 and Supplementary Figure S4). The comprehensive docking results highlight several compounds with superior binding affinities and diverse interaction profiles against the MKK3-MYC complex. Notably, the consistent involvement of Asp135 across multiple compounds underscores its pivotal role in ligand recognition. The identified compounds, particularly 4292-0516, 4476-2273, and 4476-2669, exhibit promising characteristics for further exploration as potential inhibitors or modulators of MKK3-MYC interactions.

**Figure 7.**
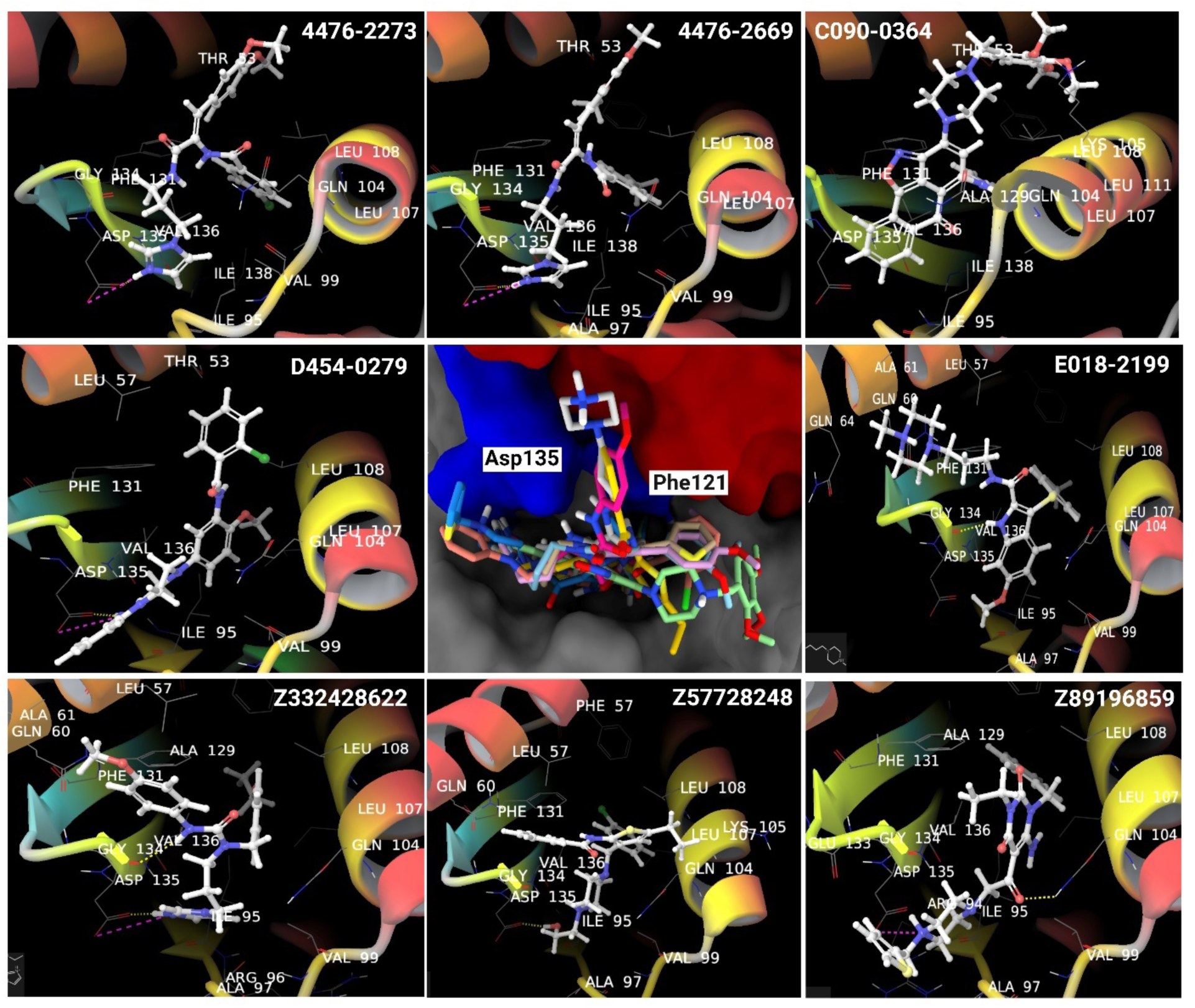
Binding poses of top lead molecules in the binding pocket of MKK3-MYC complex during docking simulation. Residues Phe121 and Asp135 show the interface region of MKK3 and MYC.

To determine the structural stability and dynamic behavior of the identified molecules within the binding pocket of the MKK3-MYC HLH LZ complex under physiological conditions, comprehensive MD simulations were conducted. Following the initial docking studies of 16,766 compounds, the top-100 compounds, chosen based on their docking scores, were used for short MD simulations (10 ns). Subsequently, a more in-depth exploration of the ligand-protein dynamics was performed through long MD simulations lasting 100 ns with three replicates. The selection of top-performing candidates for the extended MD simulations was guided by the average MM/GBSA scores, ultimately choosing the top-11 molecules. (Table 5)

**Table 5.** MM/GBSA scores (kcal/mol), standard deviation and anti-QSAR activity of top hit molecules selected from ChemDiv and Enamine.

| ID Name | $\Delta G$ 1<sup>st</sup> run | SD 1<sup>st</sup> run | $\Delta G$ 2<sup>nd</sup> run | SD 2<sup>nd</sup> run | $\Delta G$ 3<sup>rd</sup> run | SD 3<sup>rd</sup> run | Average of 3 replicas | Average of three replicate SD | SD MM/GBSA | Cancer QSAR |
| --- | --- | --- | --- | --- | --- | --- | --- | --- | --- | --- |
| C090-0364 | -69.19 | 7.90 | -86.15 | 5.55 | -71.49 | 5.99 | **-75.61** | 6.48 | 9.19 | 0.47 |
| 4292-0516 | -70.01 | 7.45 | -60.34 | 9.26 | -61.08 | 11.10 | **-63.81** | 9.27 | 5.38 | 0.73 |
| 4476-2273 | -61.01 | 8.80 | -55.39 | 13.80 | -75.9 | 10.34 | **-64.10** | 10.98 | 10.59 | 0.74 |
| 4476-2669 | -69.58 | 6.87 | -83.21 | 9.50 | -73.55 | 7.63 | **-75.44** | 8.00 | 7.00 | 0.77 |
| D454-0279 | -80.72 | 7.57 | -55.49 | 6.33 | -77.95 | 7.05 | **-71.38** | 6.98 | 13.83 | 0.68 |
| E018-2199 | -74.37 | 6.97 | -39.74 | 14.08 | -54.54 | 7.85 | **-56.21** | 9.63 | 17.37 | 0.67 |
| Z57728248 | -61.99 | 8.93 | -74.83 | 13.08 | -69.67 | 14.31 | **-68.83** | 12.11 | 6.45 | 0.48 |
| Z89196859 | -62.82 | 5.79 | -44.6 | 14.56 | -61.86 | 5.98 | **-56.42** | 8.78 | 10.25 | 0.34 |
| Z332428622 | -96.52 | 11.10 | -72.48 | 15.88 | -72.51 | 7.18 | **-80.50** | 11.39 | 13.87 | 0.72 |
| G831-0270 | -66.31 | 6.46 | -51.52 | 6.40 | -44.76 | 11.33 | **-54.30** | 8.06 | 10.98 | 0.69 |
| G856-4409 | -72.78 | 9.05 | -54.52 | 9.34 | -46.08 | 7.45 | **-57.79** | 8.61 | 13.64 | 0.80 |
| SGI-1027 | -64.76 | 11.56 | -51.41 | 11.20 | -56.21 | 8.45 | **-57.46** | 10.40 | 6.76 | 0.55 |
| ZINC12561387 | -64.56 | 8.53 | -79.89 | 7.63 | -52.29 | 9.29 | **-65.58** | 8.49 | 13.82 | 0.71 |
| ZINC33933233 | -71.94 | 6.75 | -64.92 | 5.67 | -46.74 | 7.86 | **-61.20** | 6.76 | 13.00 | 0.74 |

These findings offer valuable insights into the molecular basis of MKK3-MYC binding and pave the way for subsequent experimental validation and drug development efforts targeting this critical signaling pathway in the context of breast cancer. Table 4 shows the top-hit candidate molecules that well fit in the interface of the MKK3-MYC complex.

### 3.8 Trajectory analysis of top hits selected from ChemDiv and Enamine

The analysis of interaction profiles was complemented by the utilization of Whisker plots, providing a comprehensive visualization of the dynamic behavior of key interactions throughout the 100 ns simulations. Whisker plots offer an insightful representation of the variability and distribution of interaction frequencies, allowing us to distinguish patterns and fluctuations in the ligand-protein interactions. The majority of molecules in the dataset exhibit low variability, with a compact interquartile range and a lack of outliers, suggesting a homogeneous and consistent distribution within the dataset (Figure 8).

**Figure 8.**
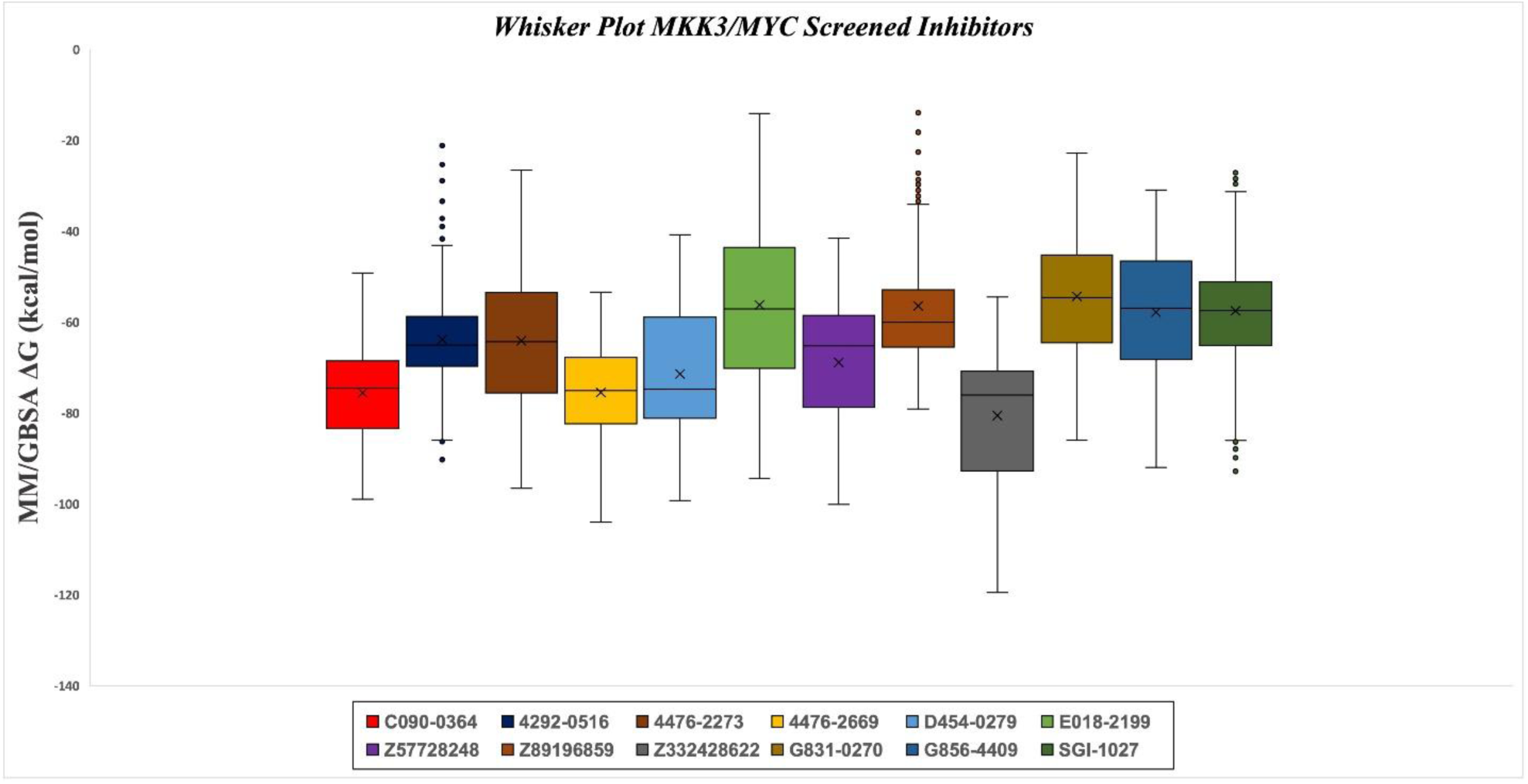
Whisker plot representation of MM/GBSA scores of identified 11 final hit compounds. The majority of molecules in the dataset exhibit low variability, with a compact interquartile range and a lack of outliers, suggesting a homogeneous and consistent distribution within the dataset.

The ensuing analysis of MD trajectories involved analyzing essential parameters, including protein’s backbone RMSD, ligand RMSD, and the characterization of interactions with crucial residues. These analyses provide a refined understanding of the ligand-induced alterations in the MKK3-MYC HLH LZ complex, offering valuable insights into the dynamic interchange between the ligands and the target protein at a molecular level. The average protein-RMSD values for the selected molecules were determined to assess their structural stability during MD simulations. Z332428622 exhibited an average RMSD of 4.62 Å, indicating moderate deviations from the initial protein structure. 4476-2273 demonstrated a lower average RMSD of 3.14 Å, suggesting a relatively stable conformation throughout the simulation. Other compounds, such as 4292-0516, 4476-2669, D454-0279, E018-2199, G831-0270, and G856-4409, displayed average RMSD values ranging from 4.25 Å to 5.43 Å. The reference molecule SGI-1027 exhibited an average RMSD of 5.40 Å. These findings provide insights into the dynamic behavior of the selected molecules, with lower RMSD values indicating more structurally stable interactions with the protein over the course of the simulations (Figure S5).

The ligand RMSD’s based on protein structure (i.e., Lig-fit-protein) RMSD averages for the selected molecules were calculated to evaluate the stability of ligand-protein complexes during the simulation. Particularly, Z332428622 exhibited a relatively low average RMSD of 3.02 Å, suggesting a stable binding conformation. In contrast, 4292-0516, 4476-2273, and E018-2199 displayed intermediate average RMSD values of 6.40 Å, 3.94 Å, and 4.88 Å, respectively. 4476-2669 and D454-0279 showed higher average RMSDs of 8.11 Å and 7.08 Å, indicating some degree of flexibility in the ligand-protein interactions. G831-0270 and G856-4409 displayed relatively elevated average RMSDs of 10.44 Å and 6.28 Å, respectively, suggesting more dynamic binding behaviors. The reference molecule, SGI-1027, exhibited a substantial average RMSD of 10.54 Å. These results provide insights into the varying degrees of stability and flexibility in the ligand-protein complexes, crucial for understanding their dynamic behavior during the simulation time (Figure S6).

Identified compounds maintain significant interactions with the interface region of MKK3 and MYC HLH LZ domain. Among the top-ranked compounds, Z332428622 is the top candidate that shows an average MM/GBSA score of -80.50 kcal/mol and well fit in the binding pocket of the MKK3-MYC HLH LZ complex during the 100 ns MD simulations. Hydrophobic contacts were established with key residues, including Ile67, Phe131, Val99, and Val136, contributing to the structural stabilization of the ligand-protein complex. Particularly, water-mediated bridges involving Asp133 and Glu133 were observed, indicating dynamic interactions that potentially modulate the binding affinity. Furthermore, a hydrogen bond was observed with Glu70 that was maintained at 28% during the 100 ns simulation time. Quantitative analysis of the interaction frequencies revealed a consistent and substantial involvement of Phe131, with a sustained 38% interaction occurrence throughout the simulation. Additionally, Asp133 and Glu133 exhibited considerable interaction frequencies of 58% and 49%, respectively, underscoring their integral roles in maintaining the stability of the ligand-protein complex (Figure 9).

**Figure 9.**
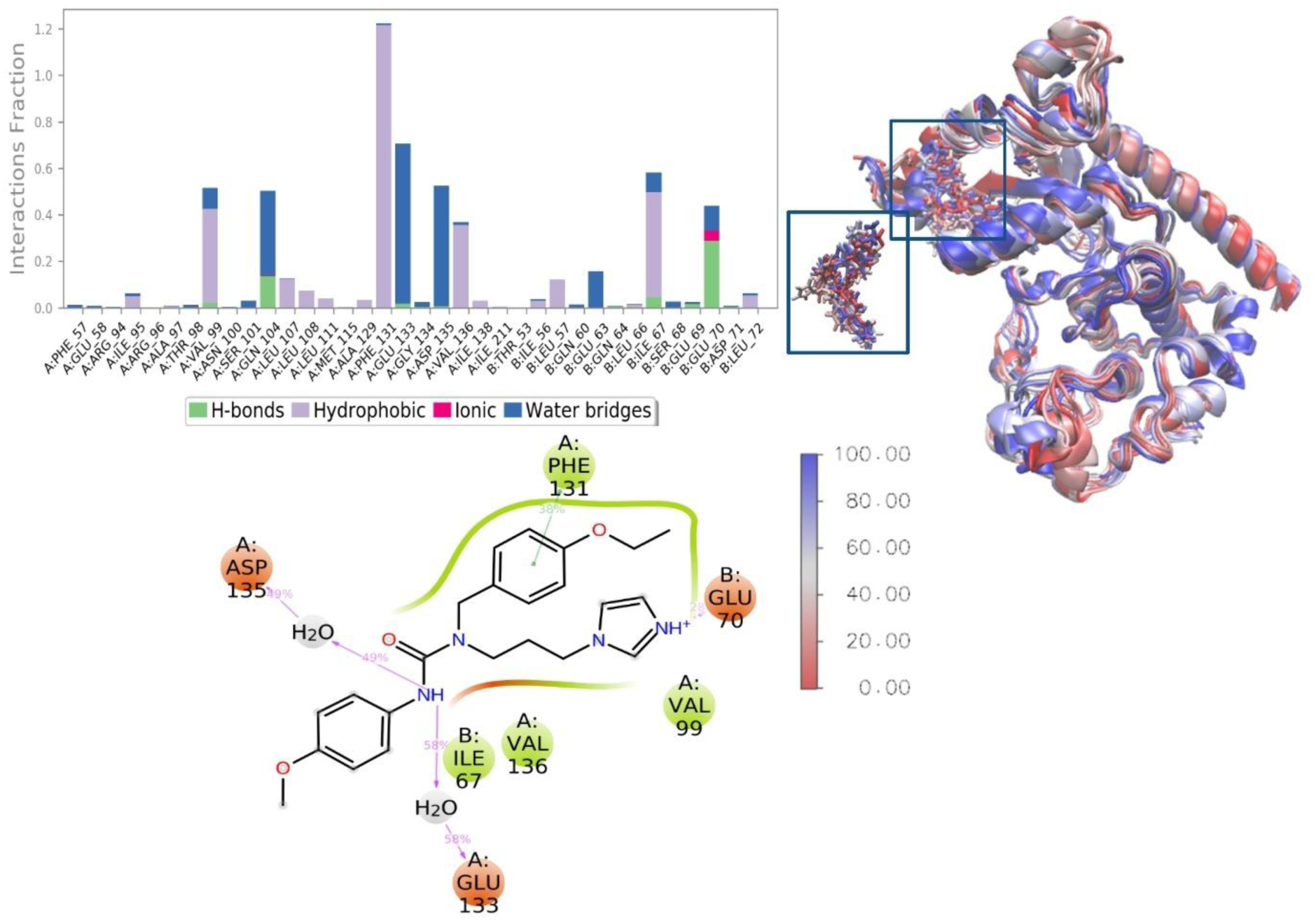
2D interaction map and change of MD poses of Z332428622 in the binding pocket of MKK3-MYC HLH LZ. Red, initial conformation; blue, final conformation.

These findings illuminate the detailed intermolecular interactions of Z332428622 within the MKK3-MYC HLH LZ binding site, offering mechanistic insights into its potential as a modulator of this crucial protein-protein interaction. Similar types of interactions are explained by Yang et al. ^10^. Simulation interactions of other hit molecules are explained in Supplementary Tables S5. Overall, these findings provide valuable insights into the intricate molecular interactions within the studied protein complexes, offering a basis for further understanding their structural and functional significance.

In the MM/GBSA analysis, SGI-1027 serves as the reference molecule for assessing the binding free energies of other compounds. The ΔG values represent the calculated binding free energies for each compound across three independent runs. The analysis provides insights into the stability of the ligand-protein complexes and the relative strength of their interactions. Observing the results, it is evident that Z332428622 exhibits the most favorable binding energy (average of three replicate simulations -80.50 kcal/mol), implying strong and stable interactions with the target protein. On the other hand, G831-0270 has the highest ΔG among the ligands (average of three replicate simulations, -54.30 kcal/mol), indicating a relatively weaker binding affinity compared to other ligands. The standard deviation (SD) associated with each set of runs provides insights into the consistency and reliability of the calculated binding free energies. Lower SD values suggest more reliable predictions, while higher values may indicate variability or uncertainty in the results. It’s important to note that the average of the three replica runs (Average_3replica) provides a consolidated view of the overall binding affinity for each compound. Therefore, compounds with lower average ΔG values are considered to have more stable and stronger binding interactions with the target protein. MM/GBSA analysis highlights the binding affinities of various compounds relative to the reference molecule SGI-1027. The results suggest potential lead compounds with favorable binding energies that could be further explored in drug development efforts (Table 5).

### 3.9 MetaCore/MetaDrug Analysis

Identified 11 compounds were also used in binary QSAR analysis. For this aim, we used MetaCore/MetaDrug platform ^41^ of Clarivate Analytics. Cancer-QSAR model was used for the therapeutic activity predictions. This functionality facilitates the evaluation of a compound’s potential for anticancer activity prediction. During the screening process, the input compounds undergo comparison with those demonstrating established high anticancer activity. The Cancer-QSAR model is employed to predict therapeutic activity values (TAV) for the compounds, with TAV values normalized within the range of 0 to 1. A TAV exceeding 0.5 suggests potential anticancer properties. The cutoff value of 0.5 was established. Since three molecules, namely C090-0364, Z57728248, and Z89196859 had smaller than 0.5 TAV, they were excluded, and only those with a predicted cancer TAV of at least 0.5 were chosen for subsequent analysis. (Table 5)

The top 8 molecules again were subjected to a Swiss similarity search. 8 cycles were run and each cycle we got 400 molecules that were used for molecular docking and MD simulations. Among the analogs, only 2 molecules (ZINC12561387 and ZINC33933233) demonstrated enhanced binding affinity compared to initial reference compounds. While ZINC12561387 has a docking score of -8.34 kcal/mol and an average MM/GBSA score of -73.63 kcal/mol, ZINC33933233 with a docking score of -8.17 kcal/mol and average MM/GBSA score of -69.15 kcal/mol (Table 5 and Figures S5 and S6).

### 3.10 Steered MD Simulations

When examining the structural and functional characteristics of a receptor-ligand complex, sMD simulations are a powerful tool that can reveal important stabilizing elements as well as the receptor’s mechanical flexibility. An external force is applied along a designated direction vector to start the unbinding of the ligand from the receptor. sMD simulations were used to pull the ligand away from the binding pocket of the protein at a constant rate of 0.1 Å/ps, aimed away from the binding interface, to assess the binding affinity of hit compounds to the target protein.

Two representative frames (i.e., the lowest and the second RMSDs to the average structure during 100 ns MD simulations) were taken from each of the three replicas used in the MD simulations for this experiment to determine the average backbone RMSD values of the hit compounds. Thus, each compound is represented by a total of six frames. An automated sMD script is developed by our group (https://github.com/DurdagiLab) that is used to pull compounds one at a time to do the sMD simulations for each frame. A 500 ps pull simulation with spring constant of 250 kj/mol.nm^2^ is applied. To determine which compounds are the most stable and require greater force to disrupt their connections with the protein, a force-time plot was produced, as illustrated in (Figure 10). This figure shows the force variation over 500 ps pull MD simulations. Compounds that exhibit higher forces may indicate greater stability within the binding pocket, necessitating more force to disrupt their interactions with the target protein. Each plot in Figure 10 represents the change in force observed between consecutive frames from the replica MD simulations of various compounds. These simulations are based on data derived from two specific frames selected from the three replicas. The figures illustrate that the compounds G856-4409, Z332428622, 4476-2669, and ZINC12561387, which exhibit greater stability, require additional force.

**Figure 10.**
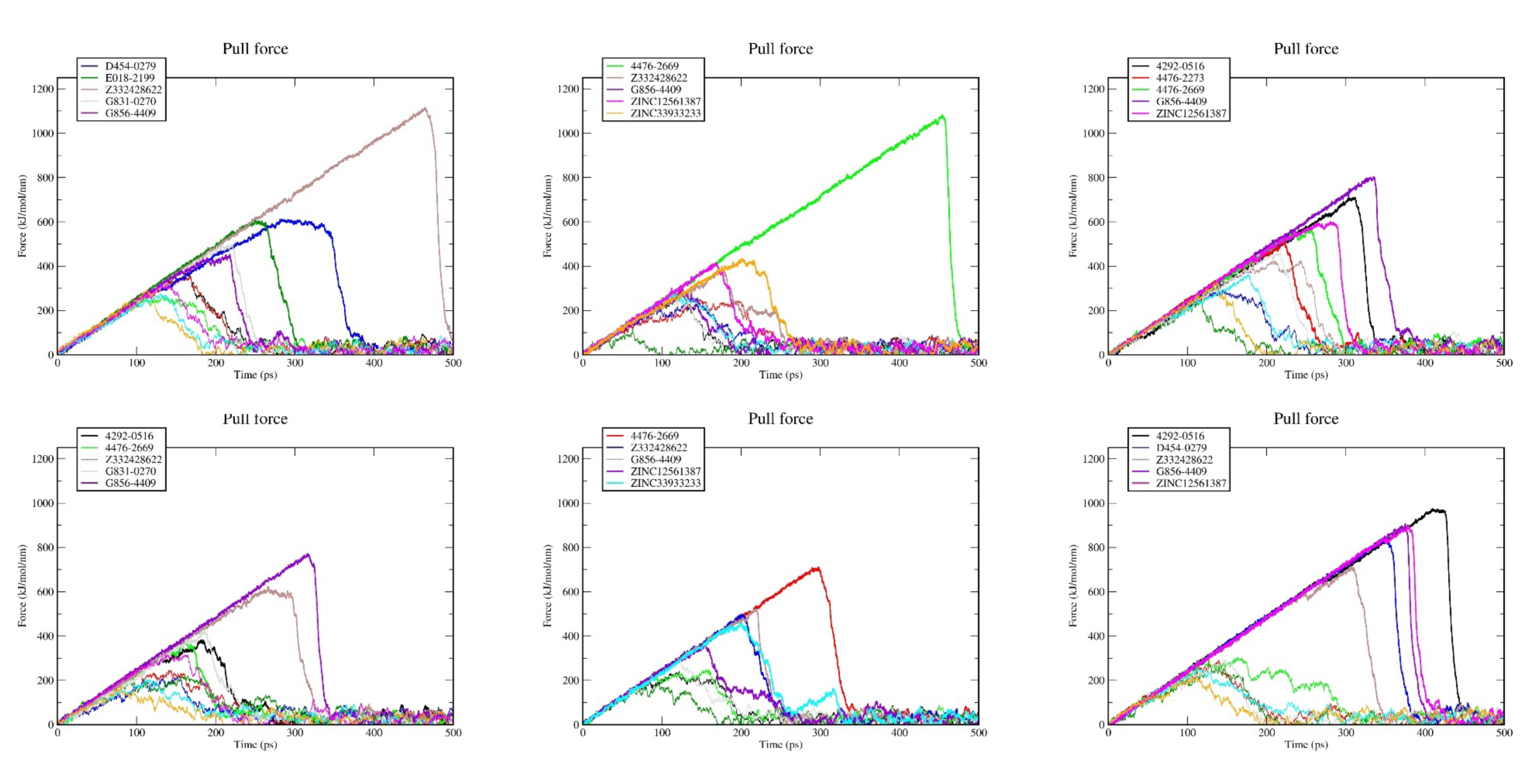
The frames with the lowest RMSD in each of the three replicas are arranged from left to right on top, and the 2^nd^ lowest RMSD frames in each of the three replicas are arranged from left to right, respectively, in bottom. This illustrates the force change that takes place during the 500 ps pulling simulations.

There is a correlation observed between MM/GBSA scores and the maximum force needed to disrupt the binding pocket, as depicted in Figure 11. This correlation is analyzed specifically between frames with the lowest average backbone RMSD values and the frames with the second lowest average backbone RMSD, excluding the outlier frame G856-4409.

**Figure 11.**
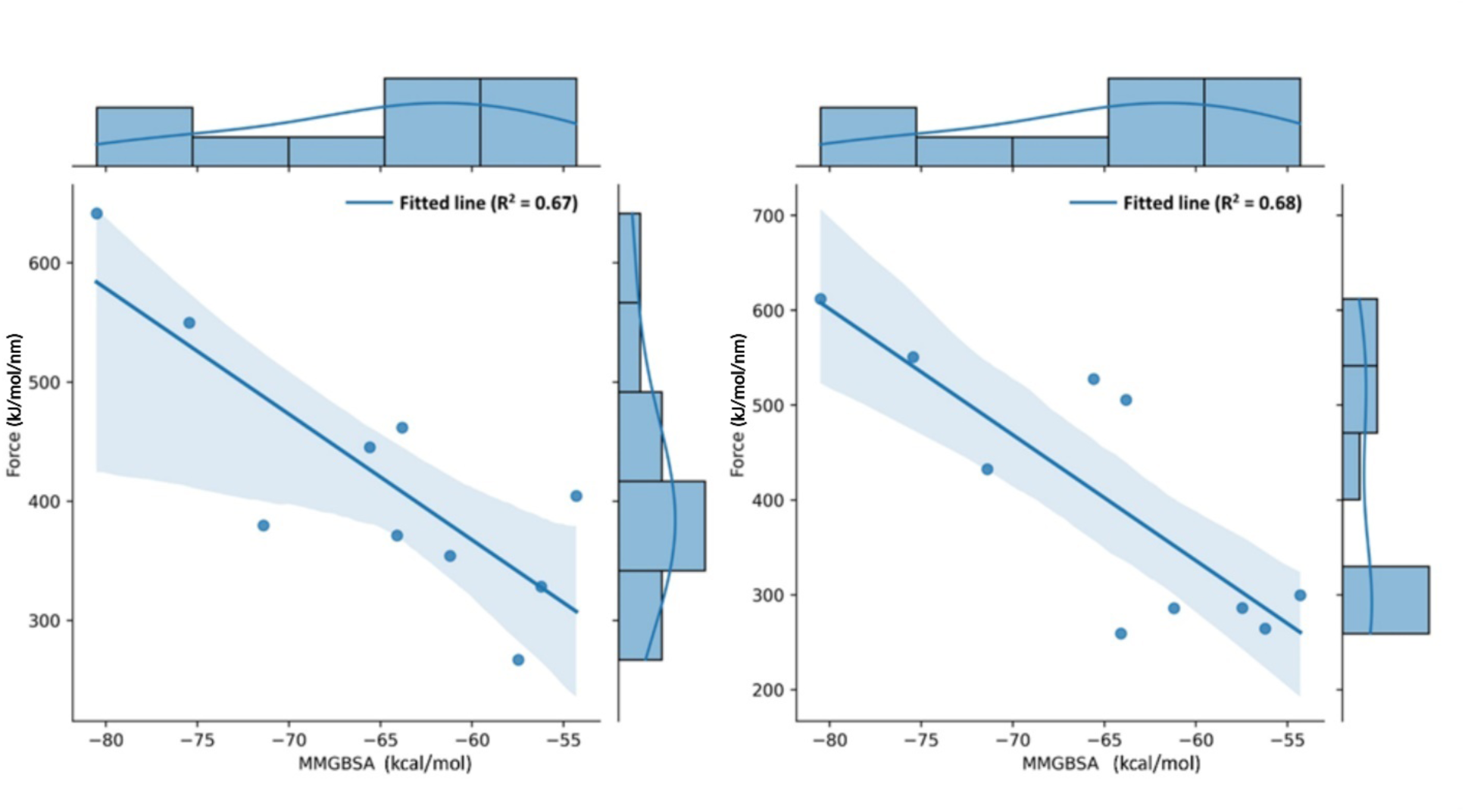
The relationship between average MMGBSA scores and forces among the lowest RMSD frame in three replicas is represented by correlation analysis on the left side. The focus of the analysis shifts to 2^nd^ lowest RMSD frame on the right side after the lowest RMSD frames are identified and their normal distribution is pointed out.

### 3.11 De novo Design of Small Molecules: Fragment-based design

Recent studies have highlighted the importance of targeting MKK3-MYC interactions in breast cancer, particularly in TNBC-related MYC activation ^42^. These interactions play a crucial role in promoting the growth and progression of TNBC, making them an attractive therapeutic target. The development of small-molecule modulators specifically designed to disrupt MKK3-MYC interactions holds great promise for the treatment of TNBC and inhibition of MYC-driven tumor growth. These small-molecule modulators could potentially have a significant impact on improving the outcomes for patients with TNBC by suppressing MYC activation and inhibiting tumor cell proliferation and metastasis ^10^. The combination of immunotherapy and targeted therapy has shown promising results in the treatment of advanced melanoma, with a 3-year survival rate of 63%. However, the efficacy of immunotherapy is still limited by side effects, resistance development, and a limited response rate ^43^. Therefore, exploring alternative strategies such as targeting MKK3-MYC interactions in breast cancer could provide a valuable addition to the current treatment options for TNBC, potentially improving patient outcomes and expanding the repertoire of targeted therapies for this aggressive subtype of breast cancer. By targeting MKK3-MYC interactions in triple-negative breast cancer, small-molecule modulators could potentially improve outcomes for patients by suppressing MYC activation and inhibiting tumor growth and metastasis.

After identifying promising molecules, we further subjected them to Auto Core Fragment *in silico* Screening (ACFIS) using the ACFIS2 server (http://chemyang.ccnu.edu.cn/ccb/server/ACFIS2/) to explore the potential for designing novel compounds. The newly designed molecules were then employed for docking studies. However, despite this effort, we did not observe any new molecules that exhibited superior docking scores compared to the initially identified compounds. This outcome suggests our initial screening approach and the efficacy of the identified molecules in the context of their binding affinity within the MKK3-MYC interaction interface (Figure S7).

### 3.12 Effect of identified small molecules to disrupt interactions between MKK3 and MYC

Interaction energies between MKK3 and MYC were measured in the absence of any hit ligand (apo state) at the interaction surface. Our goal was to assess the impact of our identified hit compounds on the interaction site. Average MM/GBSA scoring was conducted for five hit compounds across three replicas. Z-scores were calculated to analyze the differences in stability and binding affinity among the molecules, and a probability density graph was generated. The results evaluate the interaction stability and potential therapeutic applications of the molecular structures.The apo state complex structure (i.e., interaction energy between MKK3 and MYC) showed the lowest energy score, indicating the highest stability with the MM/GBSA score of - 206.41 kcal/mol. When the SGI-1027(control), 4476-2699, Z332428622, and ZINC12561387 molecules were present in the binding pocket, the interaction energy between MKK3 and MYC decreased and their average MM/GBSA scores were measured as -191.78, -193.62, -184.64, - 186.31 kcal/mol respectively. (Figure 12 and Figure S8) This suggests that these compounds weakened the interaction energies between the proteins, making them promising candidates for inhibitors of this PPI.

**Figure 12.**
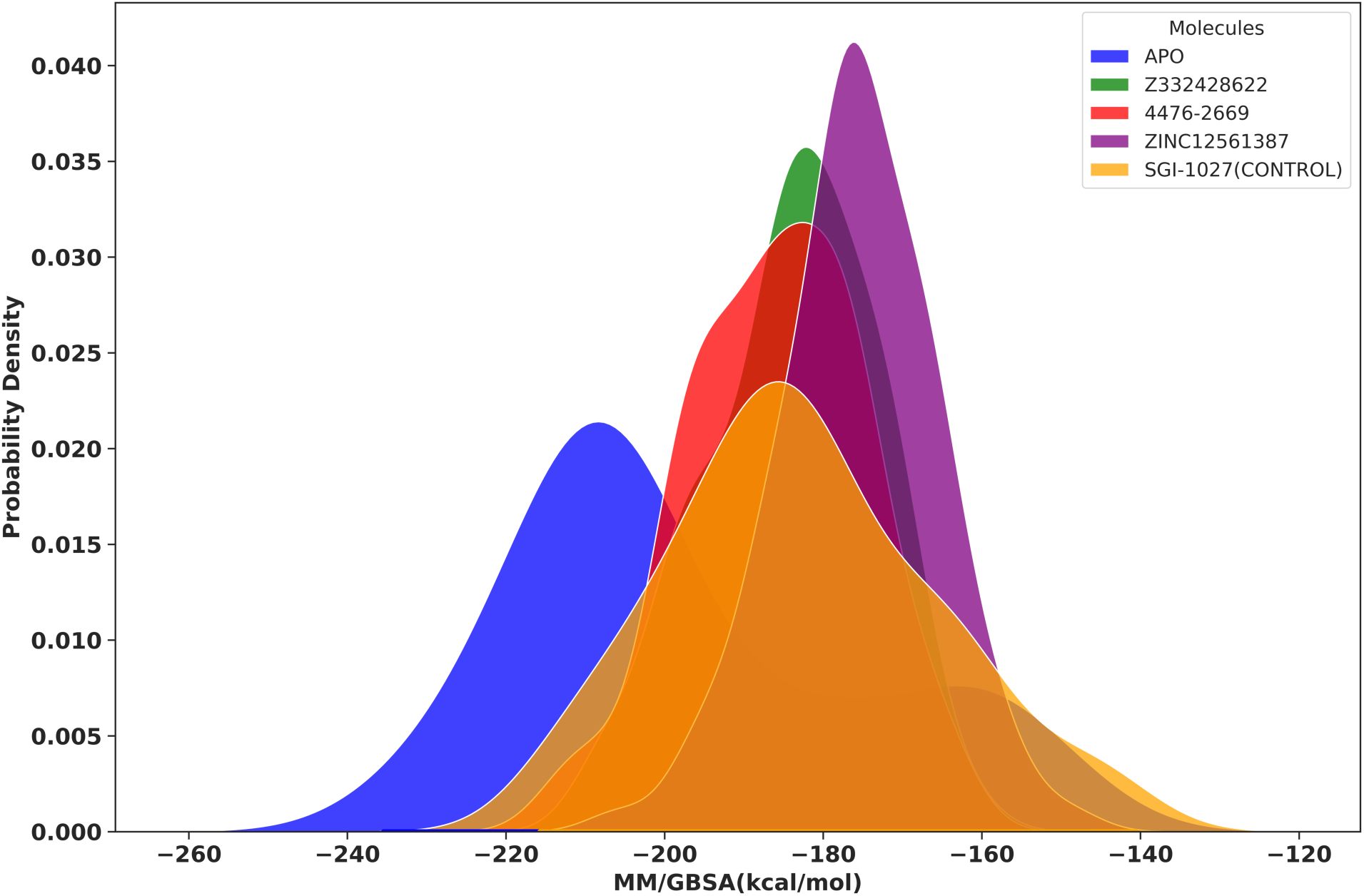
Probability densities of MM/GBSA scores for five different molecules. Blue represents the apo molecule (MKK3-MYC) without any ligand at the interaction site; green, red, purple, and orange represent Z332428622, 4476-2669, ZINC12561387, SGI-1027, respectively. Three replicas were conducted for each molecule, and the results were compared based on their energy profiles, providing information about the binding affinity and stability of the molecular structures.

## 4 Conclusions

The transcription factor MYC, a crucial regulator in cancer, represents a challenging yet critical target for therapeutic intervention. Despite the absence of FDA-approved MYC inhibitors, we address this unmet need by focusing on the inhibition of MKK3-MYC protein-protein interactions by using computer-aided drug design. Our findings show that molecules Z332428622, 4476-2669, ZINC12561387 stand out as potential lead compounds for further investigation. These findings open new avenues for drug development targeting MKK3-MYC HLH-LZ, with the identified molecules serving as promising candidates for experimental validation and potential therapeutic interventions.

## Acknowledgment

The content of this publication, prepared within the scope of the TR10/21/YEP/0133 project, which is supported by Istanbul Development Agency.

## References

(1) Bianchini, G.; Balko, J. M.; Mayer, I. A.; Sanders, M. E.; Gianni, L. Triple-Negative Breast Cancer: Challenges and Opportunities of a Heterogeneous Disease. Nat Rev Clin Oncol 2016, 13 (11), 674–690. 10.1038/NRCLINONC.2016.66.

(2) Ensenyat-Mendez, M.; Llinàs-Arias, P.; Orozco, J. I. J.; Íñiguez-Muñoz, S.; Salomon, M. P.; Sesé, B.; DiNome, M. L.; Marzese, D. M. Current Triple-Negative Breast Cancer Subtypes: Dissecting the Most Aggressive Form of Breast Cancer. Front Oncol 2021, 11. 10.3389/FONC.2021.681476.

(3) Yao, H.; He, G.; Yan, S.; Chen, C.; Song, L.; Rosol, T. J.; Deng, X. Triple-Negative Breast Cancer: Is There a Treatment on the Horizon? Oncotarget 2017, 8 (1), 1913–1924. 10.18632/ONCOTARGET.12284.

(4) Bharaj, U. K.; Lohmann, A. E.; Blanchette, P. S. Triple Negative Breast Cancer: Emerging Light on the Horizon—a Narrative Review. Precis Cancer Med 2021, 4 (0). 10.21037/PCM-20-75.

(5) Nedeljković, M.; Damjanović, A. Mechanisms of Chemotherapy Resistance in Triple-Negative Breast Cancer—How We Can Rise to the Challenge. Cells 2019, Vol. 8, Page 957 2019, 8 (9), 957. 10.3390/CELLS8090957.

(6) Chapdelaine, A. G.; Sun, G. Challenges and Opportunities in Developing Targeted Therapies for Triple Negative Breast Cancer. Biomolecules 2023, Vol. 13, Page 1207 2023, 13 (8), 1207. 10.3390/BIOM13081207.

(7) Boyle, P. Triple-Negative Breast Cancer: Epidemiological Considerations and Recommendations. Ann Oncol 2012, 23 *Suppl 6* (SUPPL. 6). 10.1093/ANNONC/MDS187.

(8) Zajac, K. K.; Malla, S.; Babu, R. J.; Raman, D.; Tiwari, A. K. Ethnic Disparities in the Immune Microenvironment of Triple Negative Breast Cancer and Its Role in Therapeutic Outcomes. Cancer Rep (Hoboken) 2023, 6 *Suppl 1* (Suppl 1). 10.1002/CNR2.1779.

(9) Shakhpazyan, N. K.; Mikhaleva, L. M.; Bedzhanyan, A. L.; Sadykhov, N. K.; Midiber, K. Y.; Konyukova, A. K.; Kontorschikov, A. S.; Maslenkina, K. S.; Orekhov, A. N. Long Non-Coding RNAs in Colorectal Cancer: Navigating the Intersections of Immunity, Intercellular Communication, and Therapeutic Potential. Biomedicines 2023, Vol. 11, Page 2411 2023, 11 (9), 2411. 10.3390/BIOMEDICINES11092411.

(10) Yang, X.; Fan, D.; Troha, A. H.; Ahn, H. M.; Qian, K.; Liang, B.; Du, Y.; Fu, H.; Ivanov, A. A. Discovery of the First Chemical Tools to Regulate MKK3-Mediated MYC Activation in Cancer. Bioorg Med Chem 2021, *45*. 10.1016/J.BMC.2021.116324.

(11) Xu, J.; Chen, Y.; Olopade, O. I. MYC and Breast Cancer. Genes Cancer 2010, 1 (6), 629–640. 10.1177/1947601910378691.

(12) Muhle-Goll, C.; Nilges, M.; Pastore, A. The Leucine Zippers of the HLH-LZ Proteins Max and c-Myc Preferentially Form Heterodimers. Biochemistry 1995, 34 (41), 13554–13564. 10.1021/BI00041A035/ASSET/BI00041A035.FP.PNG_V03.

(13) Ahmadi, S. E.; Rahimi, S.; Zarandi, B.; Chegeni, R.; Safa, M. MYC: A Multipurpose Oncogene with Prognostic and Therapeutic Implications in Blood Malignancies. J Hematol Oncol 2021, 14 (1). 10.1186/S13045-021-01111-4.

(14) Ivanov, A. A.; Gonzalez-Pecchi, V.; Khuri, L. F.; Niu, Q.; Wang, Y.; Xu, Y.; Bai, Y.; Mo, X.; Prochownik, E. V.; Johns, M. A.; Du, Y.; Khuri, F. R.; Fu, H. OncoPPi-Informed Discovery of Mitogen-Activated Protein Kinase Kinase 3 as a Novel Binding Partner of c-Myc. Oncogene 2017, 36 (42), 5852–5860. 10.1038/ONC.2017.180.

(15) Wallace, I. M.; Blackshields, G.; Higgins, D. G. Multiple Sequence Alignments. Curr Opin Struct Biol 2005, 15 (3), 261–266. 10.1016/J.SBI.2005.04.002.

(16) Song, Y.; Dimaio, F.; Wang, R. Y. R.; Kim, D.; Miles, C.; Brunette, T.; Thompson, J.; Baker, D. High-Resolution Comparative Modeling with RosettaCM. Structure 2013, 21 (10), 1735–1742. 10.1016/J.STR.2013.08.005.

(17) Jumper, J.; Evans, R.; Pritzel, A.; Green, T.; Figurnov, M.; Ronneberger, O.; Tunyasuvunakool, K.; Bates, R.; Žídek, A.; Potapenko, A.; Bridgland, A.; Meyer, C.; Kohl, S. A. A.; Ballard, A. J.; Cowie, A.; Romera-Paredes, B.; Nikolov, S.; Jain, R.; Adler, J.; Back, T.; Petersen, S.; Reiman, D.; Clancy, E.; Zielinski, M.; Steinegger, M.; Pacholska, M.; Berghammer, T.; Bodenstein, S.; Silver, D.; Vinyals, O.; Senior, A. W.; Kavukcuoglu, K.; Kohli, P.; Hassabis, D. Highly Accurate Protein Structure Prediction with AlphaFold. Nature 2021 596:7873 2021, 596 (7873), 583–589. 10.1038/s41586-021-03819-2.

(18) Waterhouse, A.; Bertoni, M.; Bienert, S.; Studer, G.; Tauriello, G.; Gumienny, R.; Heer, F. T.; De Beer, T. A. P.; Rempfer, C.; Bordoli, L.; Lepore, R.; Schwede, T. SWISS-MODEL: Homology Modelling of Protein Structures and Complexes. Nucleic Acids Res 2018, 46 (Web Server issue), W296. 10.1093/NAR/GKY427.

(19) Van Zundert, G. C. P.; Rodrigues, J. P. G. L. M.; Trellet, M.; Schmitz, C.; Kastritis, P. L.; Karaca, E.; Melquiond, A. S. J.; Van Dijk, M.; De Vries, S. J.; Bonvin, A. M. J. J. The HADDOCK2.2 Web Server: User-Friendly Integrative Modeling of Biomolecular Complexes. J Mol Biol 2016, 428 (4), 720–725. 10.1016/J.JMB.2015.09.014.

(20) Kaynak, B. T.; Bahar, I.; Doruker, P. Essential Site Scanning Analysis: A New Approach for Detecting Sites That Modulate the Dispersion of Protein Global Motions. Comput Struct Biotechnol J 2020, 18, 1577–1586. 10.1016/J.CSBJ.2020.06.020.

(21) Liu, Y.; Grimm, M.; Dai, W. tao; Hou, M. chun; Xiao, Z. X.; Cao, Y. CB-Dock: A Web Server for Cavity Detection-Guided Protein-Ligand Blind Docking. Acta Pharmacol Sin 2020, 41 (1), 138–144. 10.1038/S41401-019-0228-6.

(22) Bragina, M. E.; Daina, A.; Perez, M. A. S.; Michielin, O.; Zoete, V. The SwissSimilarity 2021 Web Tool: Novel Chemical Libraries and Additional Methods for an Enhanced Ligand-Based Virtual Screening Experience. International Journal of Molecular Sciences 2022, Vol. 23, Page 811 2022, 23 (2), 811. 10.3390/IJMS23020811.

(23) Harder, E.; Damm, W.; Maple, J.; Wu, C.; Reboul, M.; Xiang, J. Y.; Wang, L.; Lupyan, D.; Dahlgren, M. K.; Knight, J. L.; Kaus, J. W.; Cerutti, D. S.; Krilov, G.; Jorgensen, W. L.; Abel, R.; Friesner, R. A. OPLS3: A Force Field Providing Broad Coverage of Drug-like Small Molecules and Proteins. J Chem Theory Comput 2016, 12 (1), 281–296. 10.1021/ACS.JCTC.5B00864.

(24) Evans, D. J.; Holian, B. L. The Nose-Hoover Thermostat. J Chem Phys 1985, 83 (8), 4069–4074. 10.1063/1.449071.

(25) Martyna, G. J.; Klein, M. L.; Tuckerman, M. Nosé-Hoover Chains: The Canonical Ensemble via Continuous Dynamics. J Chem Phys 1992, 97 (4), 2635–2643. 10.1063/1.463940.

(26) Essmann, U.; Perera, L.; Berkowitz, M. L.; Darden, T.; Lee, H.; Pedersen, L. G. A Smooth Particle Mesh Ewald Method. J Chem Phys 1995, 103 (19), 8577–8593. 10.1063/1.470117.

(27) Roos, K.; Wu, C.; Damm, W.; Reboul, M.; Stevenson, J. M.; Lu, C.; Dahlgren, M. K.; Mondal, S.; Chen, W.; Wang, L.; Abel, R.; Friesner, R. A.; Harder, E. D. OPLS3e: Extending Force Field Coverage for Drug-Like Small Molecules. J Chem Theory Comput 2019, 15 (3), 1863–1874. 10.1021/ACS.JCTC.8B01026/SUPPL_FILE/CT8B01026_SI_002.ZIP.

(28) Dasmahapatra, U.; Kumar, C. K.; Das, S.; Subramanian, P. T.; Murali, P.; Isaac, A. E.; Ramanathan, K.; Balamurali, M. M.; Chanda, K. In-Silico Molecular Modelling, MM/GBSA Binding Free Energy and Molecular Dynamics Simulation Study of Novel Pyrido Fused Imidazo[4,5-c]Quinolines as Potential Anti-Tumor Agents. Front Chem 2022, 10. 10.3389/FCHEM.2022.991369/FULL.

(29) Li, J.; Abel, R.; Zhu, K.; Cao, Y.; Zhao, S.; Friesner, R. A. The VSGB 2.0 Model: A next Generation Energy Model for High Resolution Protein Structure Modeling. Proteins 2011, 79 (10), 2794–2812. 10.1002/PROT.23106.

(30) Oktay, L.; Sayyah, E.; Durdağı, S. Dynamic Structure-Based Pharmacophore Models for Virtual Screening of Small Molecule Libraries Targeting the YB-1. bioRxiv 2023, 2023.07.26.550723. 10.1101/2023.07.26.550723.

(31) Sayyah, E.; Oktay, L.; Tunc, H.; Durdagi, S. Developing Dynamic Structure-Based Pharmacophore and ML-Trained QSAR Models for the Discovery of Novel Resistance-Free RET Tyrosine Kinase Inhibitors Through Extensive MD Trajectories and NRI Analysis. ChemMedChem 2024, 19 (12), e202300644. 10.1002/CMDC.202300644.

(32) Kanan, T.; Kanan, D.; Erol, I.; Yazdi, S.; Stein, M.; Durdagi, S. Targeting the NF-ΚB/IκBα Complex via Fragment-Based E-Pharmacophore Virtual Screening and Binary QSAR Models. J Mol Graph Model 2019, 86, 264–277. 10.1016/J.JMGM.2018.09.014.

(33) Ikram, S.; Ahmad, J.; Rehman, I. U.; Durdagi, S. Potent Novel Inhibitors against Hepatitis C Virus NS3 (HCV NS3 GT-3a) Protease Domain. J Mol Graph Model 2020, 101. 10.1016/J.JMGM.2020.107727.

(34) Ikram, S.; Ahmad, F.; Ahmad, J.; Durdagi, S. Screening of Small Molecule Libraries Using Combined Text Mining, Ligand- and Target-Driven Based Approaches for Identification of Novel Granzyme H Inhibitors. J Mol Graph Model 2021, 105. 10.1016/J.JMGM.2021.107876.

(35) Shi, X. X.; Wang, Z. Z.; Wang, F.; Hao, G. F.; Yang, G. F. ACFIS 2.0: An Improved Web-Server for Fragment-Based Drug Discovery via a Dynamic Screening Strategy. Nucleic Acids Res 2023, 51 (W1), W25–W32. 10.1093/NAR/GKAD348.

(36) Jo, S.; Kim, T.; Iyer, V. G.; Im, W. CHARMM-GUI: A Web-Based Graphical User Interface for CHARMM. J Comput Chem 2008, 29 (11), 1859–1865. 10.1002/JCC.20945.

(37) Darden, T.; York, D.; Pedersen, L. Particle Mesh Ewald: An NsLog(N) Method for Ewald Sums in Large Systems. J Chem Phys 1993, 98, 5648. 10.1063/1.464397.

(38) Lalith Perera, *,‡; Thomas A. Darden, §; Robert E. Duke, ‖; Divi Venkateswarlu, ‖ and; Lee G. Pedersen*, §,‖. Early Unfolding Response of a Stable Protein Domain to Environmental Changes†. J Phys Chem A 2004, 108 (45), 9834–9840. 10.1021/JP048385L.

(39) Yang, X.; Liu, Y.; Gan, J.; Xiao, Z. X.; Cao, Y. FitDock: Protein-Ligand Docking by Template Fitting. Brief Bioinform 2022, 23 (3). 10.1093/BIB/BBAC087.

(40) Liu, Y.; Yang, X.; Gan, J.; Chen, S.; Xiao, Z. X.; Cao, Y. CB-Dock2: Improved Protein-Ligand Blind Docking by Integrating Cavity Detection, Docking and Homologous Template Fitting. Nucleic Acids Res 2022, 50 (W1), W159–W164. 10.1093/NAR/GKAC394.

(41) Ikram, S.; Ahmad, J.; Durdagi, S. Screening of FDA Approved Drugs for Finding Potential Inhibitors against Granzyme B as a Potent Drug-Repurposing Target. J Mol Graph Model 2020, 95. 10.1016/J.JMGM.2019.107462.

(42) Luo, L.; Tang, H.; Ling, L.; Li, N.; Jia, X.; Zhang, Z.; Wang, X.; Shi, L.; Yin, J.; Qiu, N.; Liu, H.; Song, Y.; Luo, K.; Li, H.; He, Z.; Zheng, G.; Xie, X. LINC01638 LncRNA Activates MTDH-Twist1 Signaling by Preventing SPOP-Mediated c-Myc Degradation in Triple-Negative Breast Cancer. Oncogene 2018, 37 (47), 6166–6179. 10.1038/S41388-018-0396-8.

(43) Haist, M.; Stege, H.; Kuske, M.; Bauer, J.; Klumpp, A.; Grabbe, S.; Bros, M. Combination of Immune-Checkpoint Inhibitors and Targeted Therapies for Melanoma Therapy: The More, the Better? Cancer Metastasis Rev 2023, 42 (2), 481–505. 10.1007/S10555-023-10097-Z.

