## supplementary files for "Employing Steered MD simulations for Effective Virtual Screening: Active Pharmacophore Search by Dynamic Corrections to target MKK3-MYC Interactions"

#### Supplementary Tables

**Table S1. (top)** 2D structure and docking score of SGI-1027. (bottom) Docking results of compounds obtained from SwissSimilarity search.

| ID | 2D Structure | Docking Score (kcal/mol) | Ligand Efficiency (kcal/mol) |
|---|---|---|---|
| ZINC95591603<br><br>(SGI-1027) | 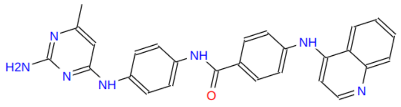 | -5.33 | −0.15<br><br>35 Heavy atoms |

| ID | 2D Structure | Docking Score (kcal/mol) | Ligand Efficiency (kcal/mol) |
|---|---|---|---|
| ZINC72442345 | 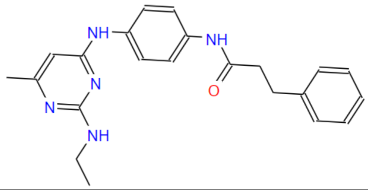 | -6.60 | −0.24<br><br>28 Heavy atoms |
| ZINC329038688 | 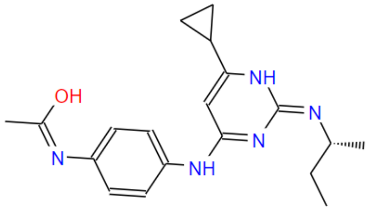 | -6.45 | −0.26<br><br>25 Heavy atoms |
| ZINC329041878 | 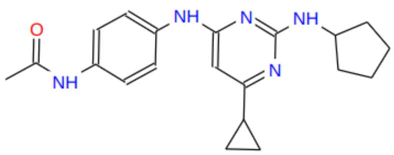 | -6.37 | −0.24<br><br>26 Heavy atoms |

| | | | |
|---|---|---|---|
| ZINC329040800 | 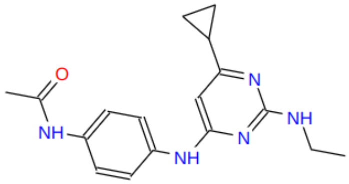 | -6.34 | −0.28<br>23 Heavy atoms |
| ZINC72442345 | 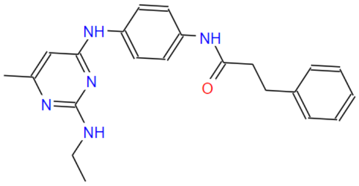 | -6.33 | −0.23<br>28 Heavy atoms |
| ZINC329066602 | 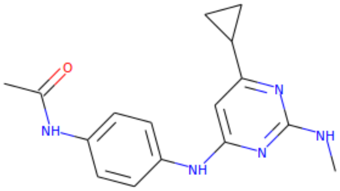 | -6.14 | −0.28<br>22 Heavy atoms |
| ZINC1579058 | 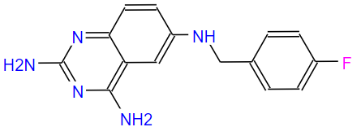 | -6.12 | −0.29<br>21 Heavy atoms |
| ZINC329014006 | 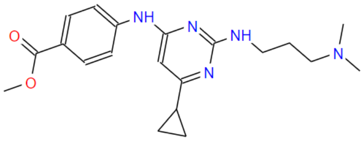 | -6.08 | −0.23<br>27 Heavy atoms |
| ZINC329038689 | 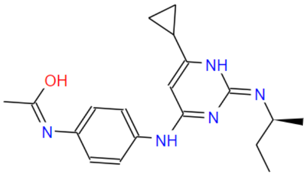 | -6.06 | −0.12<br>52 Heavy atoms |
| ZINC328956371 | 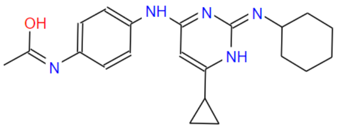 | -6.00 | −0.11<br>56 Heavy atoms |

**Table S2.** ZINC72442345 (-6.50 kcal/mol, docking score) is used for SwissSimilarity. Table shows docking results of analog compounds.

| ID | 2D Structure | Docking Score (kcal/mol) | Ligand Efficiency (kcal/mol) |
|---|---|---|---|
| ZINC329065661 | 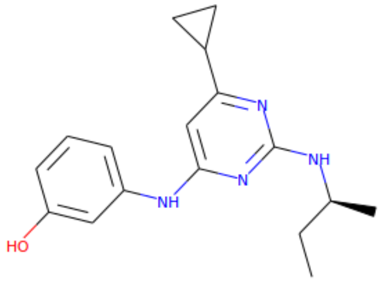 | -6.95 | −0.32<br><br>22 Heavy atoms |
| ZINC328982170 | 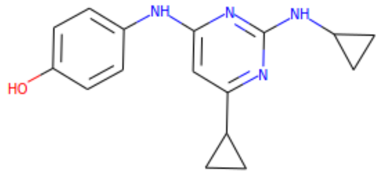 | -6.76 | −0.32<br><br>21 Heavy atoms |
| ZINC328976613 | 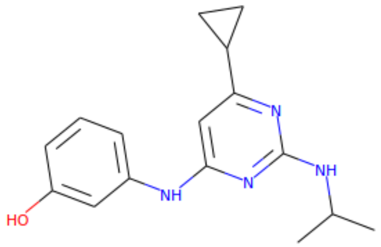 | -6.63 | −0.32<br><br>21 Heavy atoms |

| | | | |
|---|---|---|---|
| ZINC328959347 | 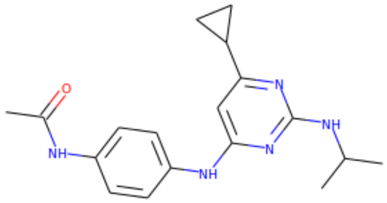 | -6.59 | −0.27<br><br>24 Heavy atoms |
| ZINC328961426 | 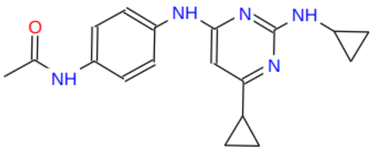 | -6.52 | −0.27<br><br>24 Heavy atoms |
| ZINC329015850 | 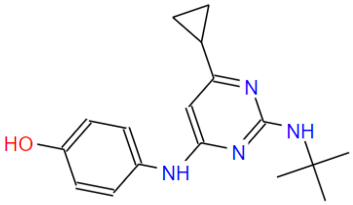 | -6.51 | −0,30<br><br>22 Heavy atoms |
| ZINC329034814 | 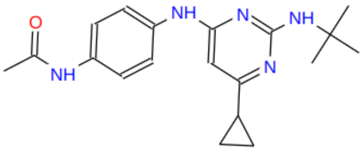 | -6.50 | −0,26<br><br>25 Heavy atoms |
| ZINC329038689 | 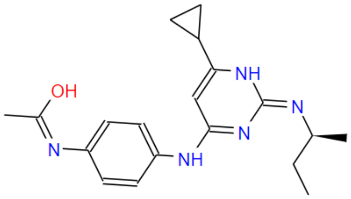 | -6.41 | −0.26<br><br>25 Heavy atoms |
| ZINC329065660 | 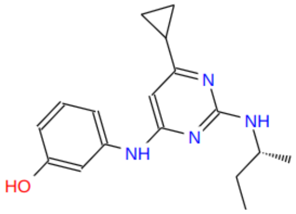 | -6.40 | −0.29<br><br>22 Heavy atoms |

| | | | |
|---|---|---|---|
| ZINC217504902 | 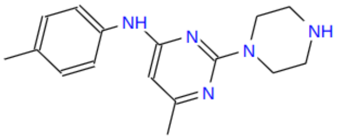 | -6.39 | −0.30<br><br>21 Heavy atoms |

**Table S3.** ZINC329001568 (-9.01 kcal/mol, docking score) is used for SwissSimilarity. Table shows docking results of analog compounds.

| ID | 2D Structure | Docking Score (kcal/mol) | Ligand Efficiency (kcal/mol) |
|---|---|---|---|
| ZINC328987638 | 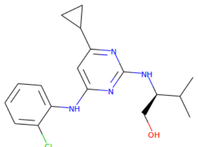 | -7.38 | −0.31<br><br>24 Heavy Atoms |
| ZINC329047693 | 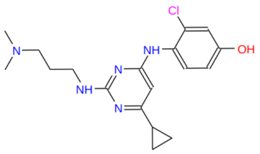 | -7.37 | −0.29<br><br>25 Heavy Atoms |
| ZINC329044265 | 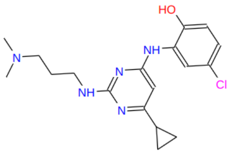 | -7.37 | −0.29<br><br>25 Heavy Atoms |
| ZINC328984509 | 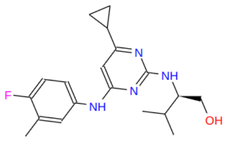 | -7.27 | −0.29<br><br>25 Heavy Atoms |
| ZINC328984510 | 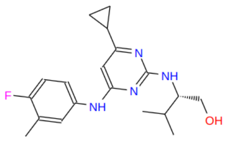 | -7.27 | −0.29<br><br>25 Heavy Atoms |
| ZINC328969042 | 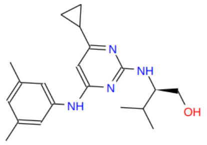 | -7.06 | −0.28<br><br>25 Heavy Atoms |

| | | | |
|---|---|---|---|
| ZINC328969037 | 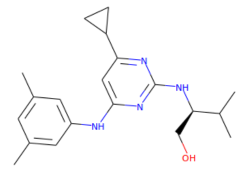 | -7.06 | 0.28<br><br>25 Heavy Atoms |
| ZINC329069759 | 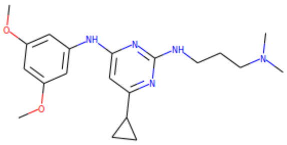 | -7.05 | −0.26<br><br>27 Heavy Atoms |

**Table S4.** ZINC328987638 (-7.38 kcal/mol, docking score) is used for SwissSimilarity. Table shows docking results of analog compounds.

| ID | 2D Structure | Docking Score (kcal/mol) | Ligand Efficiency (kcal/mol) |
|---|---|---|---|
| ZINC328974372 | 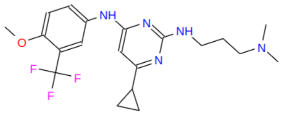 | -8.91 | −0.36<br><br>29 Heavy Atoms |
| ZINC328977658 | 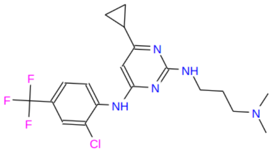 | -8.91 | −0.32<br><br>28 Heavy Atoms |
| ZINC329061341 | 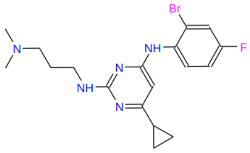 | -8.85 | −0.35<br><br>25 Heavy Atoms |
| ZINC329070534 | 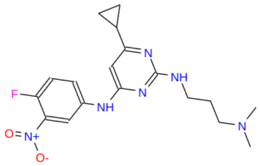 | -8.71 | −0.32<br><br>27 Heavy Atoms |
| ZINC329001473 | 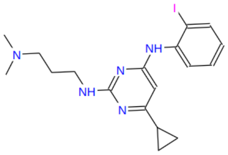 | -8.70 | −0.36<br><br>24 Heavy Atoms |

| | | | |
|---|---|---|---|
| ZINC244718869 | 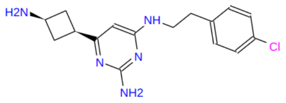 | -8.26 | −0.38<br><br>22 Heavy Atoms |
| ZINC223854430 | 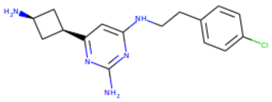 | -8.25 | −0.38<br><br>22 Heavy Atoms |
| ZINC329038614 | 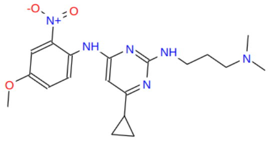 | -7.89 | −0.28<br><br>28 Heavy Atoms |
| ZINC328978812 | 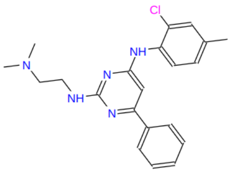 | -7.86 | −0.29<br><br>27 Heavy Atoms |

**Table S5.** Protein-ligand contact 3D pose and ligand interaction of top hits.

| Molecule ID | Docking Pose | MD simulation Ligand-Interaction | Docking Ligand Interaction |
|---|---|---|---|
| 4292-0516 | 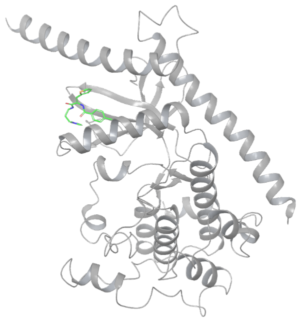 | 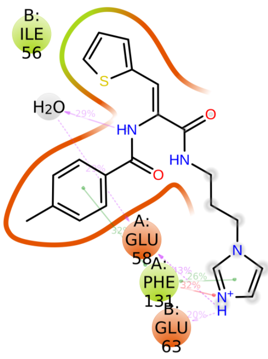  | 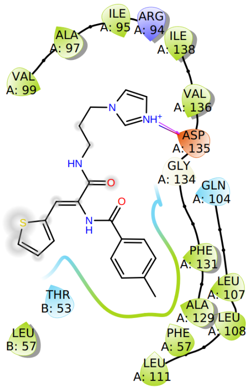  |

| | | | |
|---|---|---|---|
| 4476-2273 | 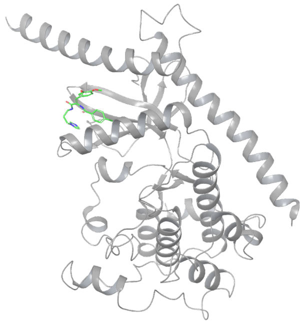 | 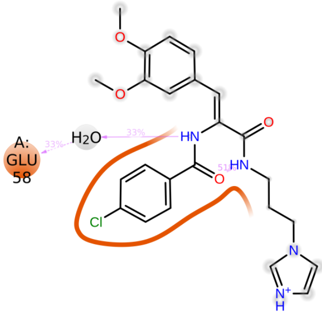 | 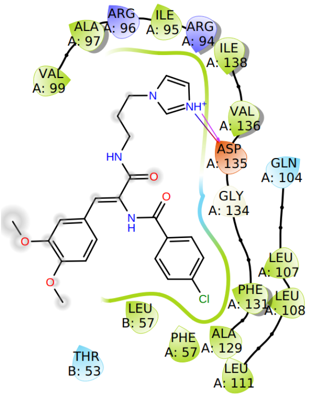 |
| 4476-2669 | 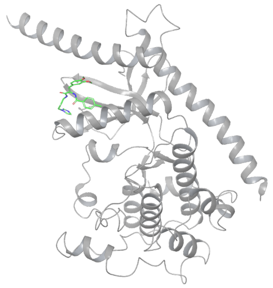 | 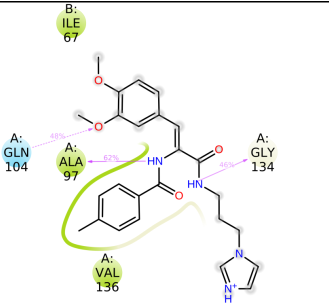 | 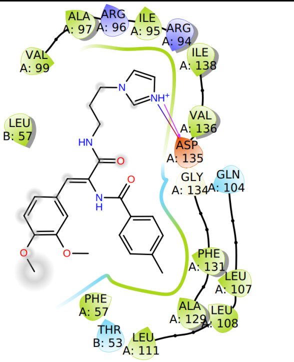 |
| D454-0279 | 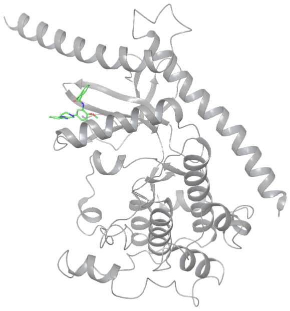 | 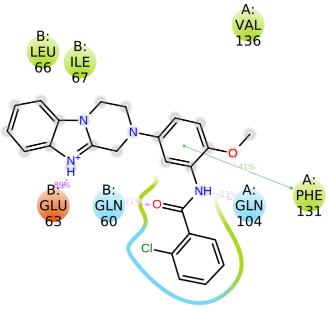 | 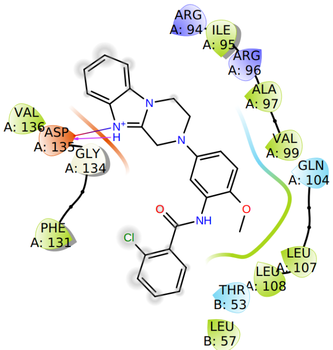 |
| E018-2199 | 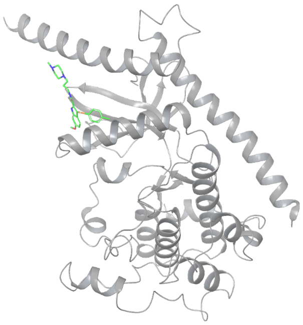 | 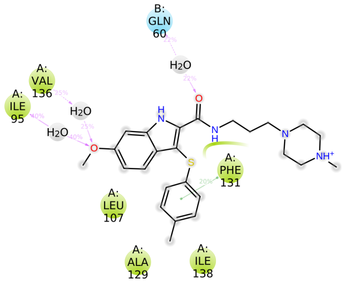 | 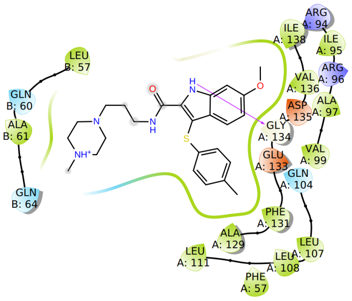 |

| | | | |
|---|---|---|---|
| G831-0270 | 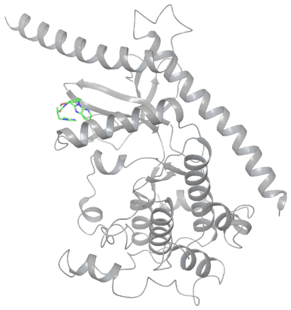 | 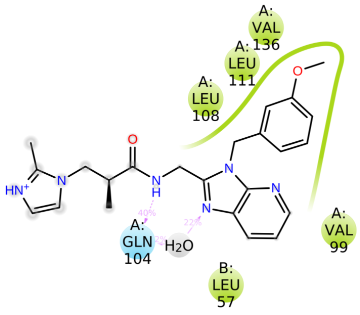 | 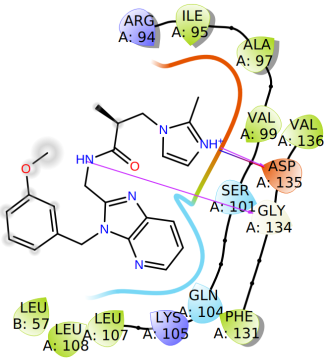 |
| G856-4409 | 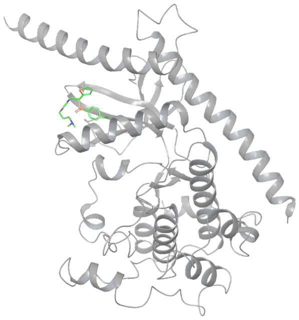 | 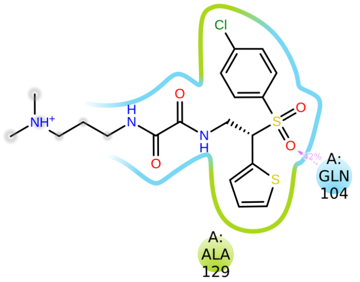 | 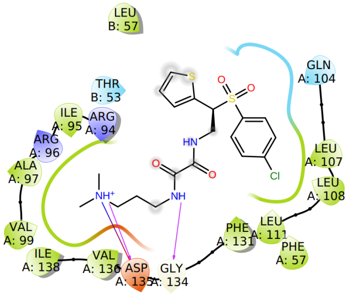 |
| SGI_1027 | 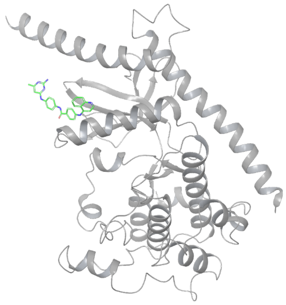 | 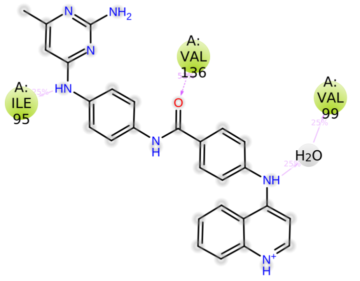 | 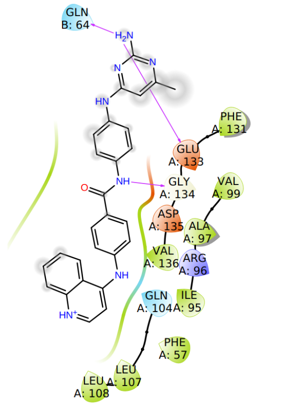 |

**Table S6.** Interactions detail of all selected hits from the pharmacophore screening.

| ID Name | 2D map | Hydrophobic | Water bridge | Hydrogen bond | Ionic interaction |
|---|---|---|---|---|---|
| 4292-0516 | 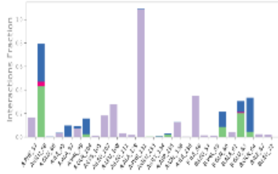 | Ile56, Phe131 | Glu58, Glu63 | Glu58 | Phe131 |

| | | | | | |
|---|---|---|---|---|---|
| 4476-2273 | 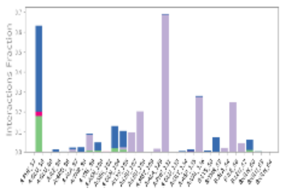 | Phe131, Val136, Ile56 | Glu58, | Glu58 | |
| 4476-2669 | 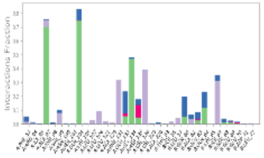 | Phe131, Val136, Ile67 | Gln104 | Ala97, Gln104, Gly134, Gln64 | Asp135 |
| D454-0279 | 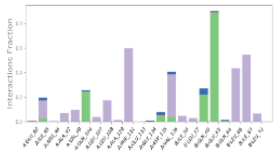 | Phe131, Val136, Leu66, Ile67 | | Glu63, Gln60 | |
| E018-2199 | 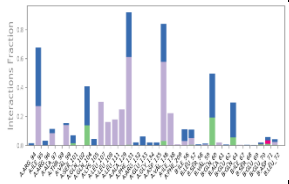 | Ile95, Leu107, Leu108, Phe131, Val136 | Ile95, Gln104 , Phe131 , Val136 , Ile138, Gln60 | Gln104, Gln60 | |
| G831-0270 | 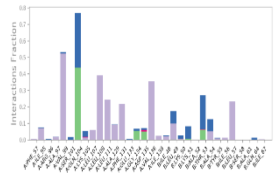 | Val99,Leu107, Leu108, Ala129, Phe131, Val136 | Gln104 , | Gln104 | Thr53 |
| G856-4409 | 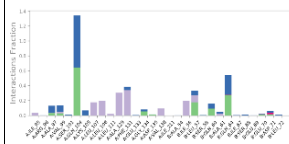 | Ala129, Phe131, | Gln104 , | Gln104,, Gln64 | |
| SGI_1027 | 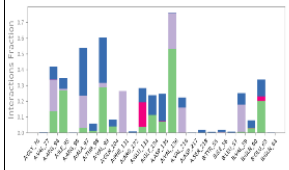 | Val77,Val97, Val136, | Val77, Val97, Val99, | Val136, Glu63, Val99, Ile95, Val77 | Glu133 |

### Supplementary Figures

a)

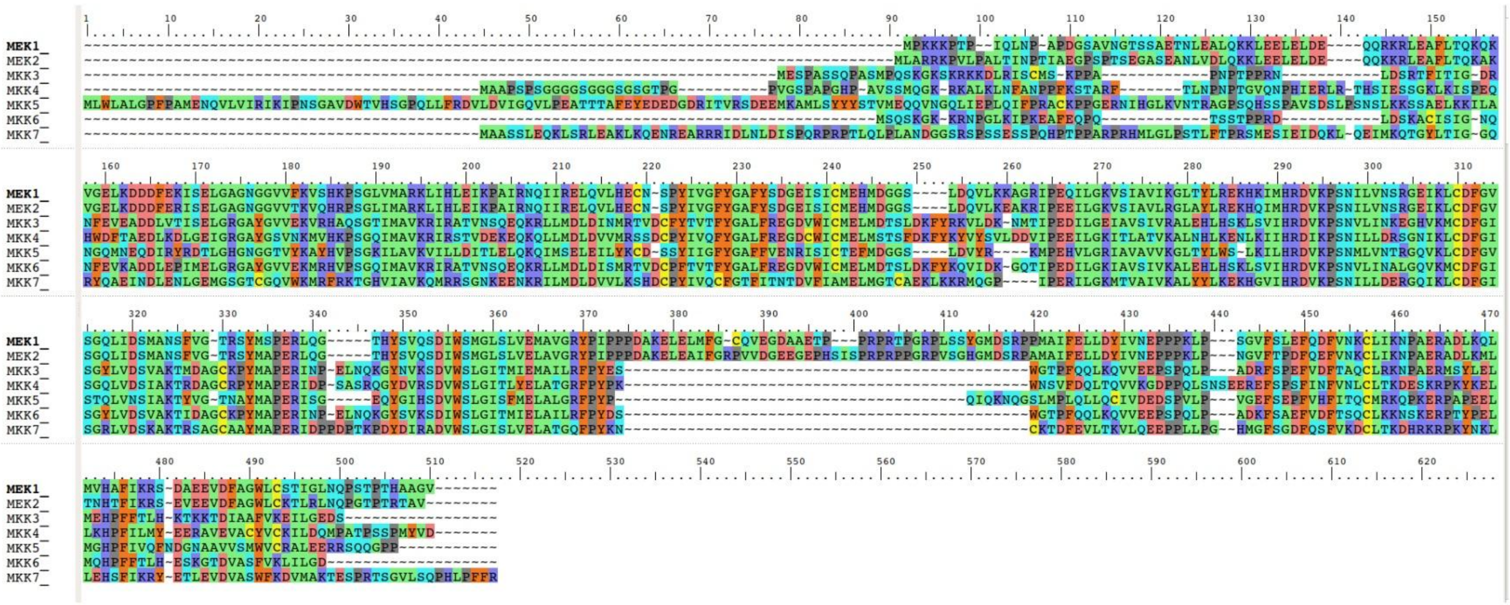

b)

| | MEK1_ | MEK2_ | MKK3_ | MKK4_ | MKK5_ | MKK6_ | MKK7_ |
|---|---|---|---|---|---|---|---|
| MEK1_ | 100 | 80 | 27 | 27 | 29 | 29 | 25 |
| MEK2_ | 80 | 100 | 27 | 27 | 28 | 29 | 26 |
| MKK3_ | 27 | 27 | 100 | 44 | 28 | 77 | 32 |
| MKK4_ | 27 | 27 | 44 | 100 | 27 | 45 | 39 |
| MKK5_ | 29 | 28 | 28 | 27 | 100 | 28 | 24 |
| MKK6_ | 29 | 29 | 77 | 45 | 28 | 100 | 33 |
| MKK7_ | 25 | 26 | 32 | 39 | 24 | 33 | 100 |

c)

| | MEK1_ | MEK2_ | MKK3_ | MKK4_ | MKK5_ | MKK6_ | MKK7_ |
|---|---|---|---|---|---|---|---|
| MEK1_ | 100 | 88 | 41 | 40 | 42 | 43 | 39 |
| MEK2_ | 88 | 100 | 41 | 40 | 42 | 44 | 39 |
| MKK3_ | 41 | 41 | 100 | 56 | 40 | 84 | 47 |
| MKK4_ | 40 | 40 | 56 | 100 | 42 | 57 | 53 |
| MKK5_ | 42 | 42 | 40 | 42 | 100 | 40 | 41 |
| MKK6_ | 43 | 44 | 84 | 57 | 40 | 100 | 47 |
| MKK7_ | 39 | 39 | 47 | 53 | 41 | 47 | 100 |

d)

| | MEK1_ | MEK2_ | MKK3_ | MKK4_ | MKK5_ | MKK6_ | MKK7_ |
|---|---|---|---|---|---|---|---|
| MEK1_ | 100 | 87 | 43 | 42 | 43 | 45 | 42 |
| MEK2_ | 87 | 100 | 43 | 41 | 43 | 44 | 42 |
| MKK3_ | 43 | 43 | 100 | 62 | 40 | 86 | 51 |
| MKK4_ | 42 | 41 | 62 | 100 | 41 | 60 | 54 |
| MKK5_ | 43 | 43 | 40 | 41 | 100 | 40 | 39 |
| MKK6_ | 45 | 44 | 86 | 60 | 40 | 100 | 50 |
| MKK7_ | 42 | 42 | 51 | 54 | 39 | 50 | 100 |

**Figure S1. (a)** Multiple sequence alignment of residues between MKK1 and MKK7 kinases. **(b)** MKK6 is the closest homolog of MKK3 sharing 77% sequence identity. **(c)** MKK6 is the closest homologue of MKK3 sharing 84% sequence homology. **(d)** MKK6 is the closest homologue of MKK3 sharing 86% sequence similarity.

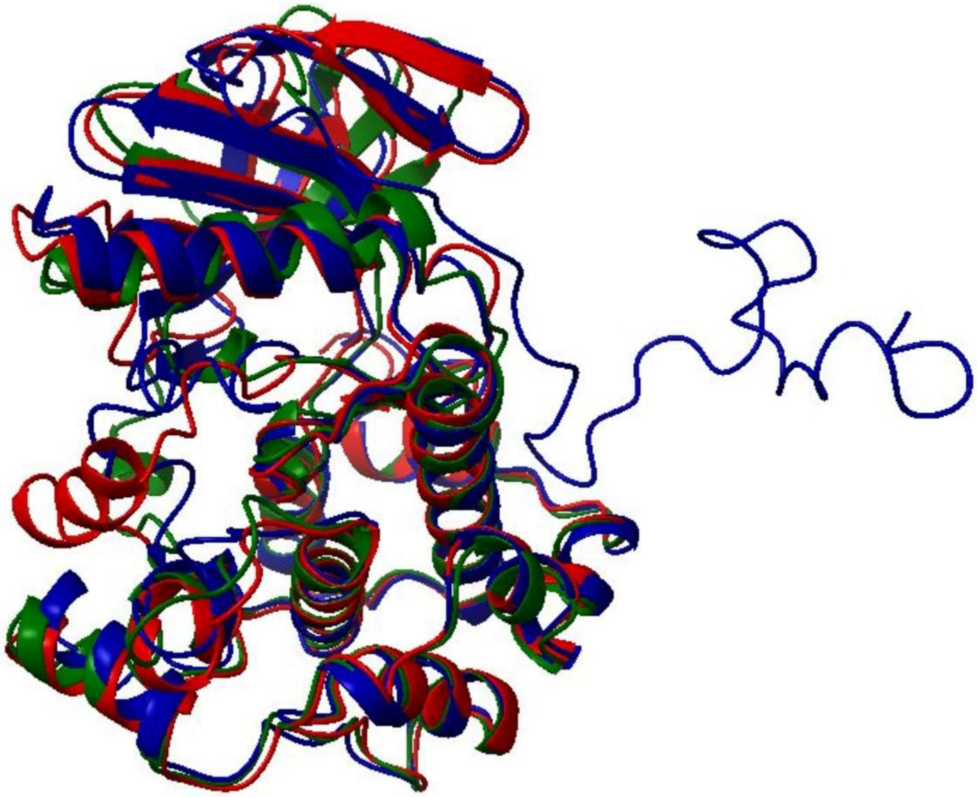

**Figure S2.** Homology model of MKK3 by using SwissModel, Rosetta, and AlphaFold. The red, blue and green colors show the model generated by AlphaFold, Rosetta, and SwissModel, respectively.

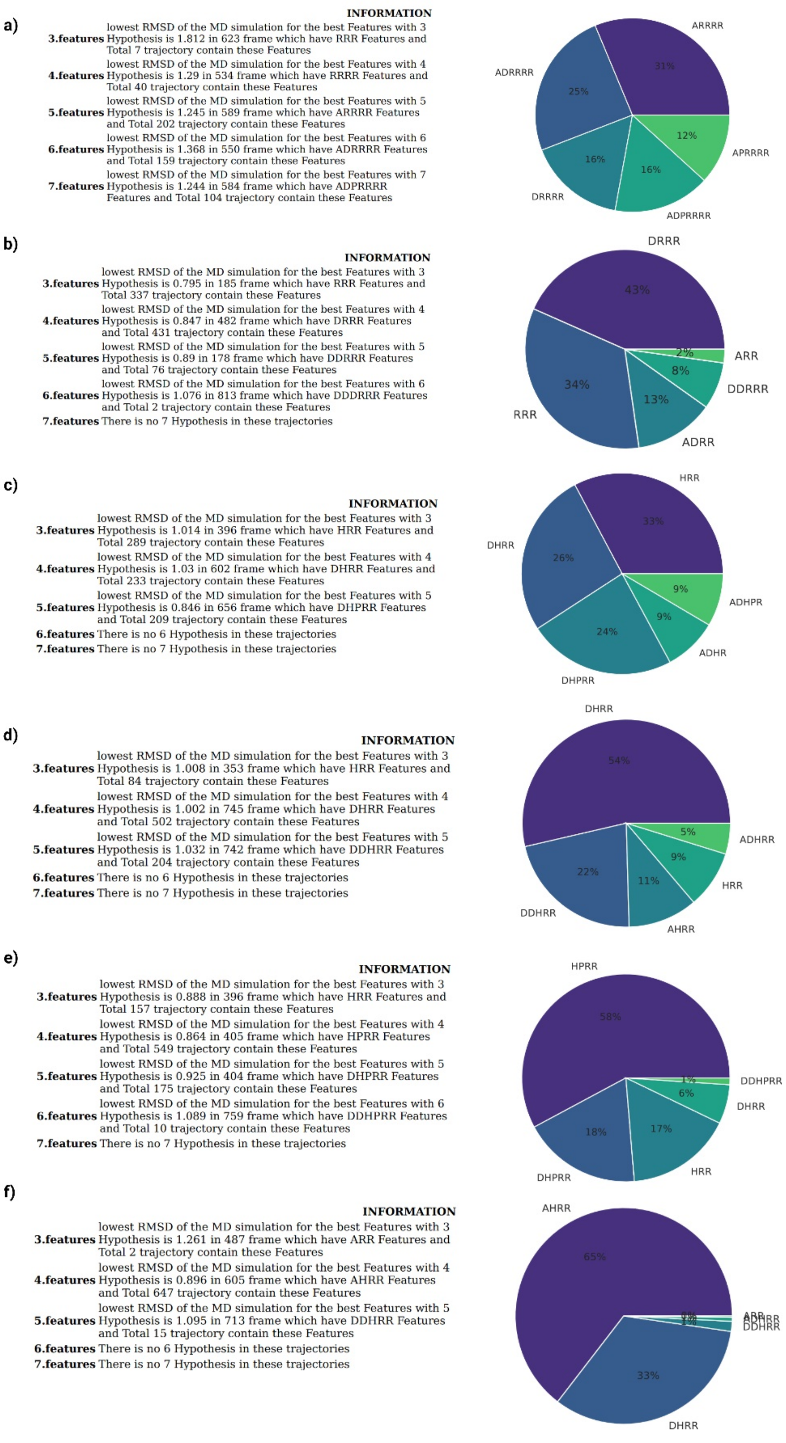


**Figure S3.** Pharmacophore feature counts and RMSD of SGI-1027 and analogs obtained from SwissSimilarity. **(a)** Represents results of SGI-1027 **(b)** Represents results of ZINC72442345 **(c)** Represents results of ZINC328974372 **(d)** Represents results of ZINC328987638 **(e)** Represents results of ZINC328991568 **(f)** Represents results of ZINC329065661.

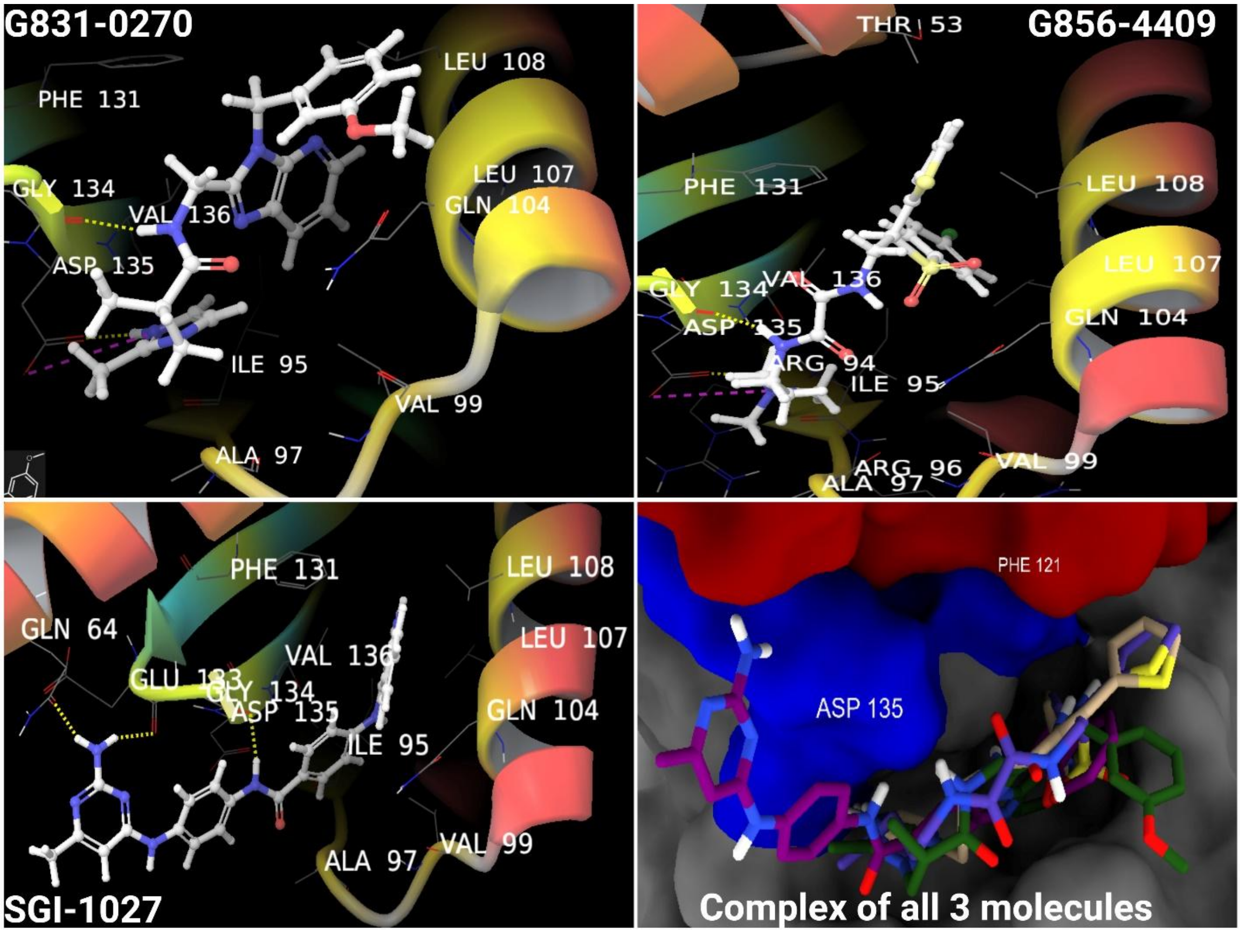


**Figure S4.** 2D and 3D interaction maps of reference molecule SGI-1027 and 2 top lead molecules in the interface region of MKK3-MYC.

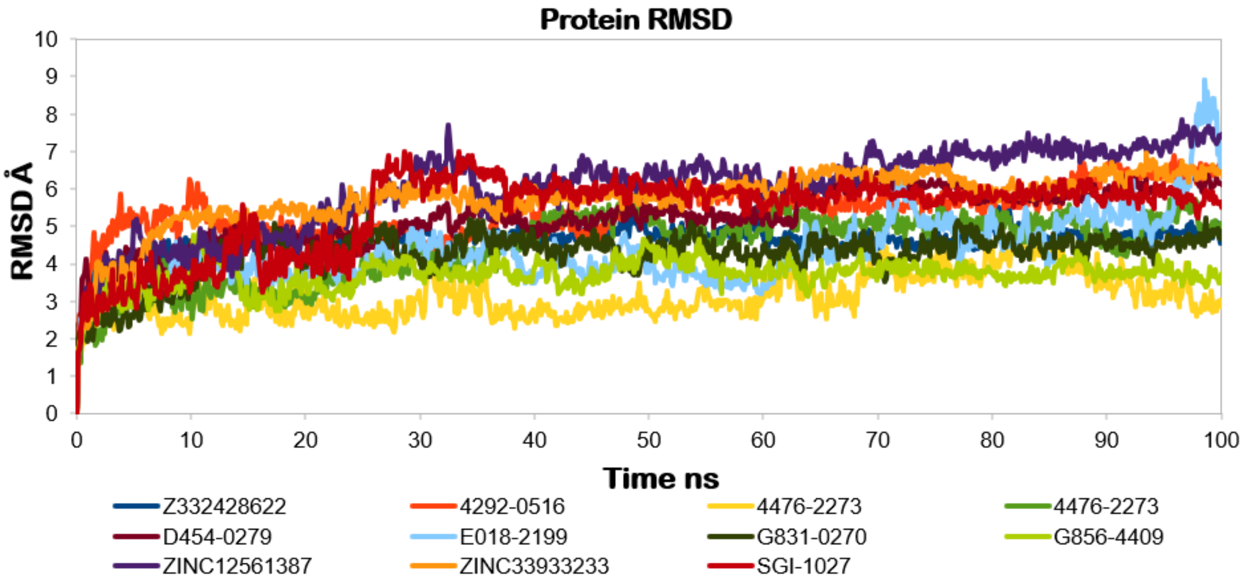


**Figure S5.** Protein backbone RMSDs of ligand-bound systems were observed during the simulation time.

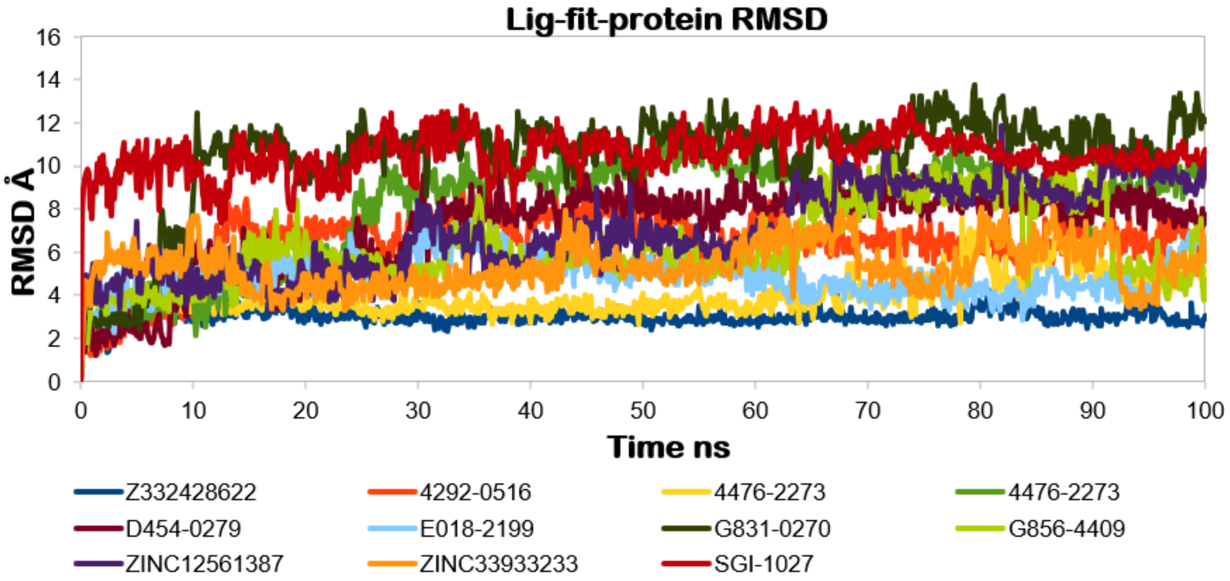


**Figure S6.** LigFitProt RMSD plots of all selected molecules from ChemDiv and enamine libraries with reported ligands SGI-1027.

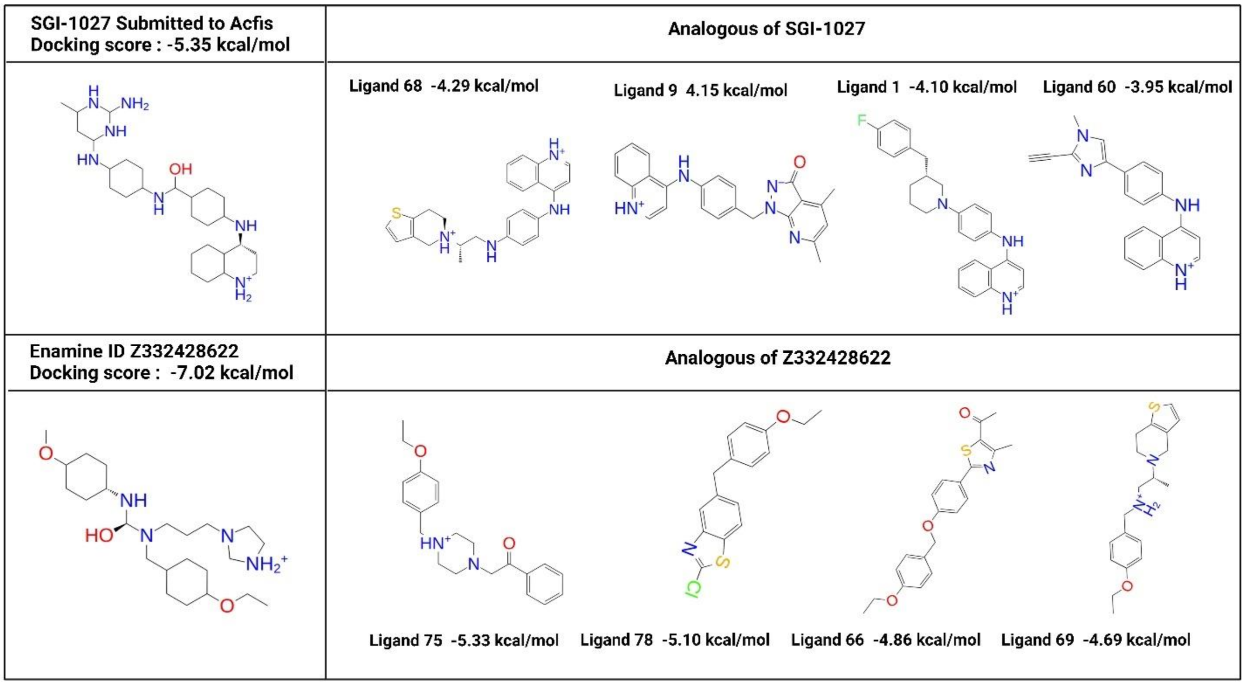


**Figure S7.** Docking score of identified molecules and 2D structure and docking score of analogous obtained from ACFIS.

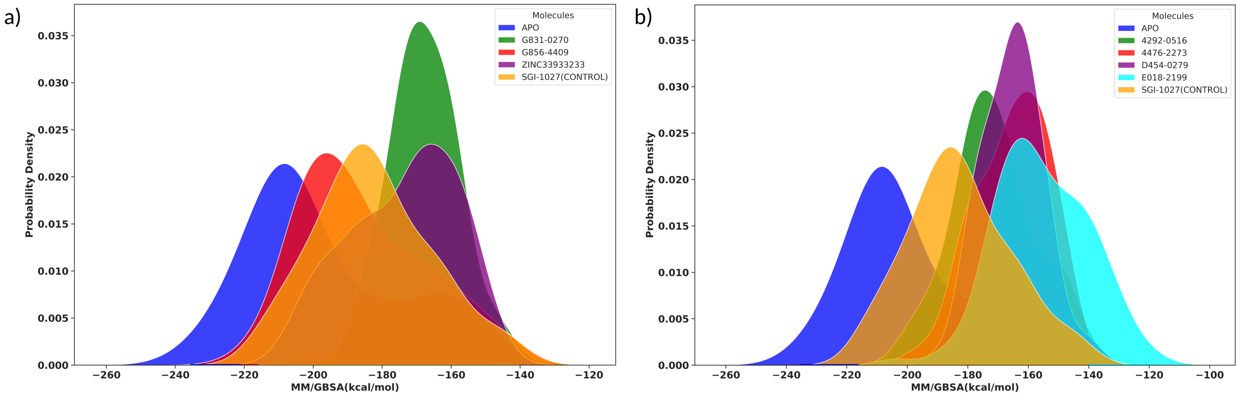


**Figure S8.** Probability densities based on the Z-scores of MM/GBSA scores for five different molecules. (a) When the lowest RMSD frame is used. Blue represents the apo state (i.e., MKK3/MYC); others represent effect of each ligand to the interaction energy between MKK3 and MYC: green, the G831-0270; red, the G856-4409; purple, ZINC33933233; and orange, the SGI-1027 control molecule. The apo state molecule exhibited the lowest energy score, indicating the highest stability, with an MM/GBSA score of -206.41 kcal/mol. When SGI-1027 (control), G831-

0270, G856-4409, and ZINC33933233 were present in the binding pocket, the interaction energy between MKK3 and MYC decreased, with MM/GBSA scores of -191.78, -182.60, -184.51, and -202.40 kcal/mol, respectively (b) When the second lowest RMSD frame is used. Blue represents the apo state; other colors represent effect of each ligand to the interaction energy between MKK3 and MYC: green, the 4292-0516; red, 4476-2273; purple, D454-0279; cyan, E018-2199; and orange, the SGI-1027 control molecule. Similarly, the apo state molecule demonstrated the lowest energy score, indicating the highest stability, with an MM/GBSA score of -206.41 kcal/mol. The presence of SGI-1027 (control), 4292-0516, 4476-2273, D454-0279, and E018-2199 in the binding pocket led to a decrease in interaction energy between MKK3 and MYC, with MM/GBSA scores of -191.78, -183.04, -185.15, -187.81, and -168.54 kcal/mol, respectively. The probability density plots show the energy profiles for each molecule, indicating their binding affinity and stability. Three repetitions were conducted for each molecule, providing comparative insights into their binding interactions.
